# Phylogenetic Synecdoche Demonstrates Optimality of Subsampling and Improves Recovery of the Blaberoidea Phylogeny

**DOI:** 10.1101/601237

**Authors:** Dominic A. Evangelista, Sabrina Simon, Megan M. Wilson, Akito Y. Kawahara, Manpreet K. Kohli, Jessica L. Ware, Benjamin Wipfler, Olivier Béthoux, Philippe Grandcolas, Frédéric Legendre

**Affiliations:** Institut de Systématique, Evolution, Biodiversité (ISYEB), Muséum national d’Histoire naturelle, CNRS, Sorbonne Université, EPHE, UA, 57 rue Cuvier, CP50, 75005 Paris, France; Department of Ecology and Evolutionary Biology, The University of Tennessee, Dabney Hall, 1416 Circle Dr., Knoxville, TN 37996, USA; Biosystematics Group, Wageningen University and Research, Droevendaalsesteeg 1, 6708 PB Wageningen, The Netherlands; Federated Department of Biological Sciences, Rutgers, The State University of New Jersey and NJIT, 195 University Ave, Newark, NJ 07102, USA; Florida Museum of Natural History, University of Florida, Gainesville, FL 32611, USA; Center for Taxonomy and Evolutionary Research, Zoological Research Museum Alexander Koenig (ZFMK), Adenauerallee 160, 53113 Bonn, Germany; CR2P (Centre de Recherche en Paléontologie – Paris), MNHN – CNRS – Sorbonne Université, UPMC Univ Paris 06, MNHN, CNRS, Paris, France; Muséum national d’Histoire naturelle, 57 rue Cuvier, CP38, 75005 Paris, France

**Keywords:** Systematics, sequence capture, targeted sequencing, anchored hybrid enrichment, Blattaria, Blattodea, Isoptera, termites

## Abstract

Phylogenomics seeks to use next-generation data to robustly infer an organism’s evolutionary history. Yet, the practical caveats of phylogenomics motivates investigation of improved efficiency, particularly when quality of phylogenies are questionable. To achieve improvements, one goal is to maintain or enhance the quality of phylogenetic inference while severely reducing dataset size. We approach this goal by designing an optimized subsample of data with an experimental design whose results are determined on the basis of phylogenetic synecdoche − a comparison of phylogenies inferred from a subsample to phylogenies inferred from the entire dataset. We examine locus mutation rate, saturation, evolutionary divergence, rate heterogeneity, selection, and *a priori* information content as traits that may determine optimality. Our controlled experimental design is based on 265 loci for 102 blaberoidean cockroaches and 22 outgroup species. High phylogenetic utility is demonstrated by loci with high mutation rate, low saturation, low sequence distance, low rate heterogeneity, and low selection. We found that some phylogenetic information content estimators may not be meaningful for assessing information content *a priori*. We use these findings to design concatenated datasets with an optimized subsample of 100 loci. The tree inferred from the optimized subsample alignment was largely identical to that inferred from all 265 loci but with less evidence of long branch attraction and improved statistical support. In sum, optimized subsampling can improve tree quality while reducing data collection costs and yielding 4-6x improvements to computation time in tree inference and bootstrapping.

## Introduction

The current “postgenomic” era of phylogenetics is characterized by large datasets and increased efforts to maximize their usage (Bravo et al. 2019 unpublished data). Yet, while more data increase potential phylogenetic information (Simon et al. 2018), they can also increase data artefacts and bias (Breinholt and Kawahara 2013; Dell’Ampio et al. 2014; Brown and Thomson 2017; Shen et al. 2017; Platt et al. 2018). In this study, we combine two approaches, data targeting and phylogenetic subsampling, aimed at improving phylogenomic inference by limiting confounding signal. Data targeting seeks to only utilize positively informative data, which requires identifying features of genetic loci that are efficient in recovering reasonable phylogenetic relationships with confidence (e.g., Townsend 2007; Borowiec et al. 2015; Gilbert et al. 2015; Tan et al. 2015; Brown and Thomson 2017; Klopfstein et al. 2017; Reddy et al. 2017; Molloy and Warnow 2018). Phylogenetic subsampling allows assessing the presence of confounding signal (Edwards 2016; Simon et al. 2018). This approach has been used to test how topologies change with taxon sampling (Evangelista et al. 2018; Simon et al. 2018), the utility of loci with different evolutionary rates (Narechania et al. 2012), the stability of nodes of biological interest (Simon et al. 2012) or the effect of missing data (Xi et al. 2016) among other phenomena (Edwards 2016).

Determining the optimality of loci (Townsend 2007; Reddy et al. 2017), as opposed to characters, must be a practical priority given that DNA sequencing technologies read contiguous regions of genomes. This is especially true when deciding which loci to target using popular genome capture strategies (e.g., Lemmon et al. 2012; Brandley et al. 2015; Gilbert et al. 2015). Although the features of individual characters are directly relevant to phylogenetic reconstruction (Dornburg et al. 2018), masking characters in alignments may still fail to improve phylogenetic accuracy (Tan et al. 2015). Here, we use the term “locus” to refer to some contiguous segment of a genome more than a few nucleotides long.

The relationship between the features and phylogenetic utility of molecular data have long been studied (e.g., see Blaxter 2004). Perhaps the prime suspect in investigating a locus’ quality for phylogenetic inference is its evolutionary rate (Klopfstein et al. 2017). The optimal mean rate of a locus for resolving difficult tree shapes (Steel and Leuenberger 2017; Dornburg et al. 2018) is very conservative (Klopfstein et al. 2017; Steel and Leuenberger 2017). Yet, targeting optimal evolutionary rates does not necessarily improve phylogenetic inference (Narechania et al. 2012; Chen et al. 2015; Doyle et al. 2015). High evolutionary rate can hinder phylogenomic studies when it leads to mutation saturation (Fong et al. 2012; Breinholt and Kawahara 2013). A locus’ evolutionary divergence among a set of taxa may better indicate its phylogenetic utility. This measure loosely accounts for phylogeny and is related to other informative features such as: long branch score (Struck 2014; Borowiec et al. 2015), evolutionary rate (Chen et al. 2015), saturation (Borowiec 2019), and entropy (Bai et al. 2013; Lewis et al. 2016). Conforming to model assumptions is another feature that should improve phylogenetic trees (Doyle et al. 2015; Reddy et al. 2017; but see Dell’Ampio et al. 2014). For instance, patterns of rate heterogeneity may be poorly estimated by the gamma distribution (Kjer and Honeycutt 2007). Finally, low selective pressure is another desirable trait. Positive selection pressure may mislead phylogenetic inference through convergent evolution, which may further induce compositional bias (Cox et al. 2014).

The most direct approach might be to measure phylogenetic information content of loci. Information content can be assessed several ways, including: (*i*) with reference to a phylogenetic tree (Townsend 2007) or (*ii*) without one (Misof et al. 2014b); (*iii*) in a Bayesian framework (Dornburg et al. 2016; Lewis et al. 2016); or (*iv*) in an empirical approach (Kück et al. 2012; Misof et al. 2014b). Using only high information content markers has sometimes improved tree congruence (Chen et al. 2015) and given greater agreement with favored morphological hypotheses (Borowiec et al. 2015). Yet, assessing information content before all data is collected (*a priori*) is challenging because new taxa can influence how character information is interpreted (Venditti et al. 2006; Hugall and Lee 2007).

We examine the effect of locus quality on empirical phylogenomic inference using a subsampling method. Subsampling to infer locus quality (Narechania et al. 2012; Edwards et al. 2017) or the reliability of data matrix features (e.g., missing data; Xi et al. 2016) have been done before but a more common usage of subsampling is to assess support for trees (e.g., Soltis and Soltis 2003; Simon et al. 2012; Edwards 2016; Evangelista et al. 2018; Simmons et al. 2018; Simon et al. 2018). We test six features suspected to affect the accuracy of phylogenetic inference, including: mutation rate, mutation saturation, pairwise sequence distance, among site rate heterogeneity, selective pressure, and *a priori* phylogenetic information content.

Phylogenetic subsampling has commonly been used to assess clade support or the effect of missing data (Edwards 2016) but it has rarely been applied to search for the most optimal tree. Despite the trend of ever-increasing dataset sizes in phylogenetics, there are yet advantages to utilizing subsamples of data (Bravo et al. 2019 unpublished data). An alignment subsample can contain a higher density of information content than whole alignments (Edwards 2016). From then on, a subsample of a dataset created for concentrating signal and excluding noise could be referred to as an optimized subsample. Such subsamples are not only improved in their data quality, but assist in improving efficiency of nearly every step of a phylogenetic pipeline.

To evaluate the optimality of a subsample of empirical data, we propose the *phylogenetic synecdoche* criterion. We define phylogenetic synecdoche as: a phylogenetic pattern inferred from a subsample that is reproducing the pattern inferred from a whole. Phylogenetic synecdoche occurs when a loss in data abundance does not result in a loss of signal, which would only be expected to occur in optimized subsamples. Of course, any two phylogenetic inferences will rarely yield the exact same tree (even if the datasets are almost identical; Shen et al. 2017), particularly for a dataset with a large number of samples. Without some guideline on an arbitrarily defined topological distance below which synecdoche is demonstrated, subsamples can be experimentally compared and ones that yield trees with lower distances to the full data tree are more optimal.

Once we have demonstrated that a subsample of data is optimized we can use that subsample to infer a new tree. We would logically assume the subsample data tree to be an improvement to the tree inferred from all the data (e.g., Cox et al. 2014; Shen et al. 2017). Now we can interpret the differences between the subsample tree and the full data tree as potential improvements instead of suboptimal, as we did before establishing locus feature optimality. Figure 1 provides a conceptual visualization of the entire workflow.

**Figure 1.**
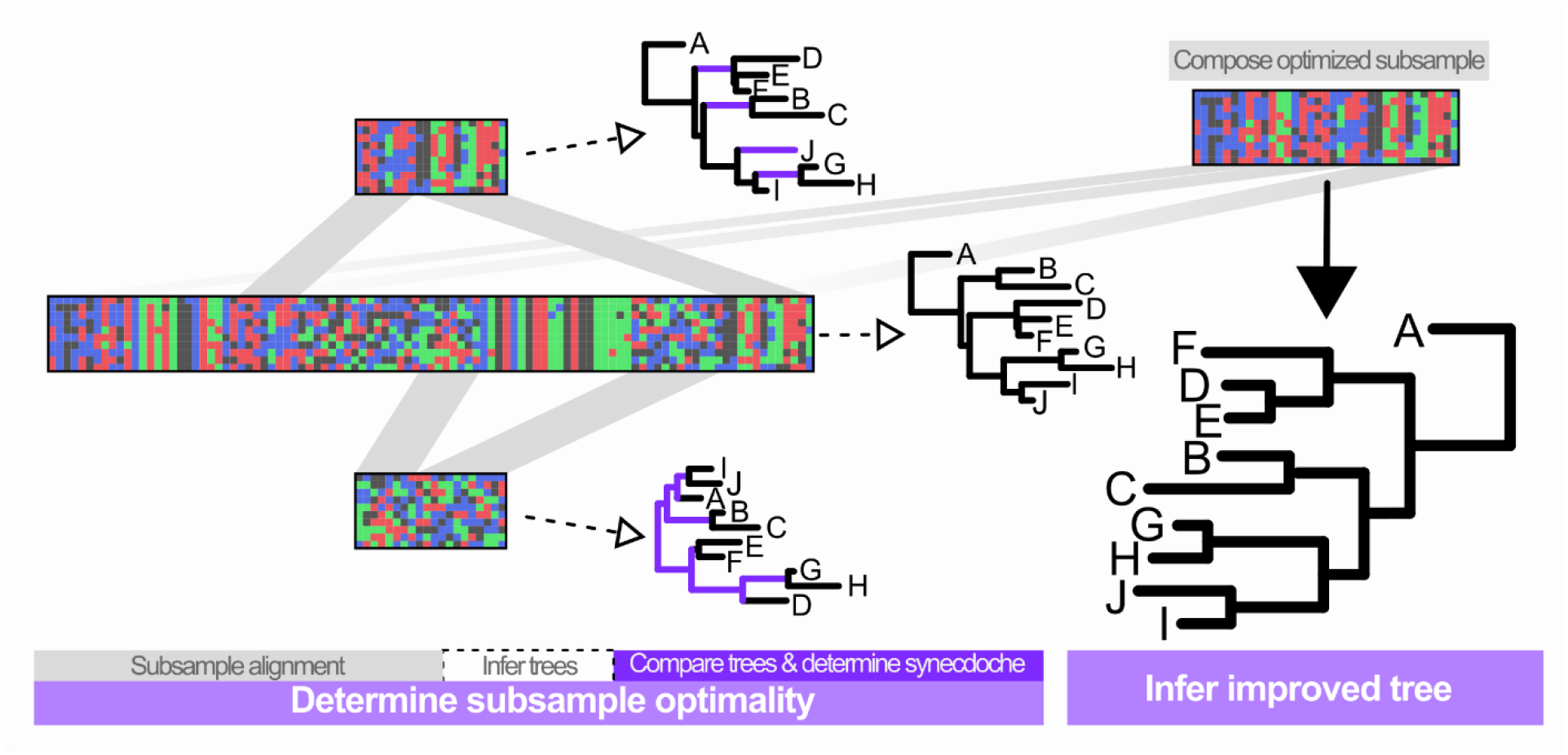
Conceptual visualization of workflow. From left to right: a large multi-locus alignment is composed; the alignment is subsampled based on features hypothesized to improve phylogenetic inference (gray); phylogenies are inferred from full and subsampled alignments (dashed arrow); resulting phylogenies are compared and the subsample’s phylogeny that has fewest differences (blue branches) from the full alignment’s phylogeny (i.e., more phylogenetic synecdoche) is deemed more optimal; all loci with the optimal trait are concatenated and a final tree is inferred.

By identifying optimized subsamples under the criterion of phylogenetic synecdoche, we attempt to recover a robust phylogeny of blaberoidean cockroaches. Blaberoidea is a 260 Myr old clade (Evangelista et al. 2019) comprising ~3600 of the ~7500 species of Blattodea (Beccaloni and Eggleton 2013). The major relationships of Blaberoidea were recently investigated in a phylogenomic study (Evangelista et al. 2019) but taxonomic sampling was limited. The major lineages of Blaberidae and many other splits occurring in the last 100 MY were not sampled (Evangelista et al. 2019). Studies that included denser taxon sampling recovered unstable topologies (Legendre et al. 2015; Legendre et al. 2017; Bourguignon et al. 2018; Evangelista et al. 2018). We combine previously published phylogenomic datasets with 265 genomic loci for 90 newly sequenced species of blaberoidean cockroaches. From the 265 loci dataset we compose two optimized subsamples of 100 loci each. By comparing the full data tree and the two subsample trees, we derive conclusions about tree plausibility and the phylogeny of Blaberoidea.

## Materials and Methods

### Taxon Sampling

We added 90 newly sequenced Blaberoidea to the Blattodea sampled in Evangelista et al. (2019) and Wipfler et al. (2019). These were chosen to cover biogeographical regions and known “tribal-level” groupings as widely as possible (supplemental data). Live samples were obtained from breeders and remaining samples were museum specimens (see NCBI accession SRP155429 for full specimen information). Tissue was taken from the middle leg of all samples. Polyneoptera outgroups were chosen among published transcriptomes (Misof et al. 2014a; Wipfler et al. 2019).

### Loci Capture and Probe Design

Using alignments of orthologous loci from Misof et al. (2014a) we extracted sequences for *Blaberus atropos* and then used BLAST+ (Camacho et al. 2009) to identify homologous sequences in transcriptomes of 17 cockroach species (*Anallacta methanoides, Asiablatta kyotensis, Blattella germanica, Cryptocercus wrighti, Diploptera punctata, Ectobius sylvestris, Ellipsidion sp., Gyna lurida, Ischnoptera deropeltiformis, Lamproblatta albipalpus, Lobopterella dimidiatipes, Nauphoeta cineria, Panchlora nivea, Paratemnopteryx couloniana, Princisia vanwaerebecki*, *Sundablatta sexpunctata*, and *Symploce* sp.). These were added to the *B. atropos* sequences and aligned with MAFFT v. 7.3 (Katoh and Standley 2013; −localpair −maxiterate 1000 −adjustdirection). The resulting 599 alignments were tested for phylogenetic information content. We define loci having high *a priori* phylogenetic information content as being able to recover a resolved topology (tree-likeness) and having high character support for non-conflicting bipartitions. First, non-tree-like data were omitted using translated amino-acid sequences analyzed in MARE v. 01.2 (Meyer et al. 2011). After having non-blaberoidean taxa removed and missing data trimmed with trimAL v. 1.2 (Capella-Gutiérrez et al. 2009), character support and conflict among nucleotide alignments was evaluated using SAMS v. 1.4.3 (Wägele and Mayer 2007). In SAMS (default options except gapmode = missing, and viewing conflicting splits with the tree) we only accepted loci for which the: (1) two most supported splits were biologically plausible (i.e. not strongly at odds with biogeographical or morphological hypotheses of relationships according to previous studies), (2) four most supported splits were not conflicting with one another, and (3) first conflicting split had less than half of the support as the highest supported split. This resulted in 92 loci with high information content. 90 additional loci that were accepted by MARE but not by the above criteria in SAMS were also chosen. Finally, 100 loci rejected by both criteria above were chosen at random.

Probes for enrichment were designed to target the final set of ~280 alignments (untrimmed, nucleotide versions) using Baitfisher v. 1.2.5 (Mayer et al. 2016). The resulting probe sequences were further quality-filtered and subsequently produced by MYcroarray.

### Library Preparation and Sequencing

In the Ware Lab at Rutgers University, Genomic DNA was extracted from 90 species (supplementary data) using the DNEasy kit (Qiagen USA) and was then sheared using a timed Fragmentase protocol and assessed with Bioanalyzer. Index-multiplexing strategies were planned and specimens were arranged on the plate to minimize the probability of cross-contaminating closely related samples. After NEBNext Ultra library preparation, indexed library pools were enriched with respect to targeted loci (MYbaits v.3.01) and then sequenced by GeneWiz USA using Illumina HiSeq’s rapid-run paired-end-250 protocol.

### Bioinformatics and Phylogenetic Inference

Demultiplexed FASTQ files had adapters removed and low-quality bases trimmed (Krueger 2017; Trimgalore v. 0.4.3 options: -q 20; -stringency 3; -e .1; −length 30). Reads were assembled with Trinity v.2.0.6 (Grabherr et al. 2011; Haas et al. 2013). Orthologs from OrthoDB v. 7 (Waterhouse et al. 2013) were identified in the 90 sequenced libraries and 37 additional Blattodea and Polyneoptera sequences (Evangelista et al. 2019) using Orthograph v. 0.6 (Petersen et al. 2017) with *Drosophila melanogaster*, *Pediculus humanus*, *Tribolium castaneum*, and *Zootermopsis nevadensis* as reference genomes [default options except --anysymbol option called in MAFFT (Katoh and Standley 2013), and blast-max-hits = 50]. 265 targeted loci were extracted from the ortholog set. Quality filtered reads are available on NCBI GenBank (SRP155429).

We aligned each locus in MAFFT v. 7.3 (Katoh and Standley 2013; options: -retree 4 -maxiterate 10 −adjustdirection) and then trimmed from the edges to eliminate leading or trailing sites missing >80% of data. A second, more thorough, alignment was conducted in MAFFT v. 7.3 (−localpair −maxiterate 1000), which was then adjusted to maintain consistent reading-frame. Alignments were finalized with manual adjustment in AliView v. 1.18 (Larsson 2014) to remove poor quality reads and correct misaligned sections. Custom scripts in Mathematica 10 (Wolfram Research, 2012) available in the package Phyloinformatica v. 0.9 (Evangelista 2019) were used to manage sequence files, translate sequences, trim and concatenate alignments. We refer to the final concatenated alignment (127 taxa) as the ”265_Full” alignment.

We ran PartitionFinder 2 (Lanfear et al. 2016) with an alignment comprising all loci and taxa. Blocks were defined as codon positions per locus and possible models were limited to GTR and GTR+G. Branch lengths were considered as unlinked, the best model was chosen with AICc, and the search scheme was rcluster (percent = 10; max = 1000). Using this codon partitioning scheme we ran a preliminary tree search with 10 independent runs in RAxML v. 8.2 (Stamatakis 2014), implemented on the CIPRES portal (Miller et al. 2010). Assessment of the best preliminary tree showed that a few taxa (*Amazonina platystylata, Doradoblatta* sp., *Ischnoptera galibi, Lanxoblatta* sp.*, Panchlora stolata, Pycnoscelus femapterus*, and *P. striata*) had exceptionally long branches. The same taxa were among those with the largest proportion of missing data (supplementary data). After reassessing the alignments in which these species were present, we removed *Pycnoscelus striata, P. femapterous*, *Ischnoptera galibi, Amazonina platystylata*, and *Doradoblatta* sp. from the analysis under the grounds that (i) their data was low quality (short reads ambiguously aligned and with many nucleotide differences) and (ii) the pattern of data presence would not allow for testing of their hypothesized taxonomic assignment (see supplementary data). When running the tree searches there were no exceedingly long branch lengths, and Blaberidae was monophyletic.

Trees were then inferred for three different alignments: 1) the full alignment (“265_Full”), 2) using only the 2^nd^ codon positions (correcting for noise; “265_2nd"), 3) low missing data alignment (correcting for relationships inferred from missing data patterns; “265_Reduced”). The latter alignment was created by only retaining nucleotide positions having data for 51 or more taxa (Phyloinformatica function trimAlign2 missingProportion = 0.60; Evangelista 2019)). The same partitioning, modelling and RAxML parameters as used above were applied to each analysis but with 100 independent tree runs (GTRGAMMA, -f d, -N 100). We also inferred one more tree using the 265_Full alignment in IQTree (Nguyen et al. 2015) using partitions determined by PartionFinder2, models determined by IQTree (Kalyaanamoorthy et al. 2017) and the options: -ninit 200 -nbest 10 -allnni -ntop 40 -wbt -wsl -wsr. These four trees (265_Full, 265_2nd, 265_Reduced, 265_Full_IQ) were later used as a baseline for establishing phylogenetic synecdoche. We assessed support for the three RAxML trees by bootstrap resampling using the auto-MRE stopping criteria (60, 300 and 108 for the three trees respectively) and calculating traditional bootstrap frequencies and node certainty scores (Kobert et al. 2016).

### Designing Test Subsamples

We calculated various statistics for each of the 265 individual alignments. Site-specific evolutionary rates were calculated using a reading-frame-adjusted concatenated alignment in fastTIGER v. 0.0.2 (Cummins and McInerney 2011; Frandsen et al. 2015). Locus mutation-saturation was calculated as the slope of a best-fit line of Hamming distances plotted against F81 corrected distances in the R package Phangorn v. 2 (Schliep 2011).

In IQ-TREE v. 1.5, mean pairwise sequence distance and non-synonymous to synonymous mutation ratio (dN/dS) were calculated under the MG model (Muse and Gaut 1994), which is known to yield less biased dN/dS values than other commonly used models (Spielman and Wilke 2015). We quantified rate heterogeneity as the number of local maxima in a frequency plot of rates after applying a Gaussian filter of scale = 25. The sum of the number of local maxima for each codon position was our estimate of rate heterogeneity, which we then length-corrected by dividing with the locus’ nucleotide length. Using Phyloinformatica v. 0.0 - 0.9 we also: counted the number of taxa per alignment, counted the proportion of ambiguous sites, measured locus length, and calculated RCFV [“relative composition of frequency variability”, a metric of nucleotide compositional bias detailed in Zhong et al. (2011)].

We then designed experimental subsamples of loci. Each subsample was designed to be non-overlapping in one of our six features (e.g., all loci with high mutation rate or all loci with low mutation rate) (Fig. 2, Table S1.2), while orthogonal to all other features (Fig. S1.1). The saturation and selection test subsamples could not be made sufficiently independent (Table S1.2), so additional controlling tests were done (see below and Fig. S4.1). A few selected loci were then added to ensure all taxa were represented in each set. *Panchlora stolata*, and *Lanxoblatta* sp, were too poorly represented among loci to include in all sets, so they were removed from all alignments and trees in our tests.

**Figure 2.**
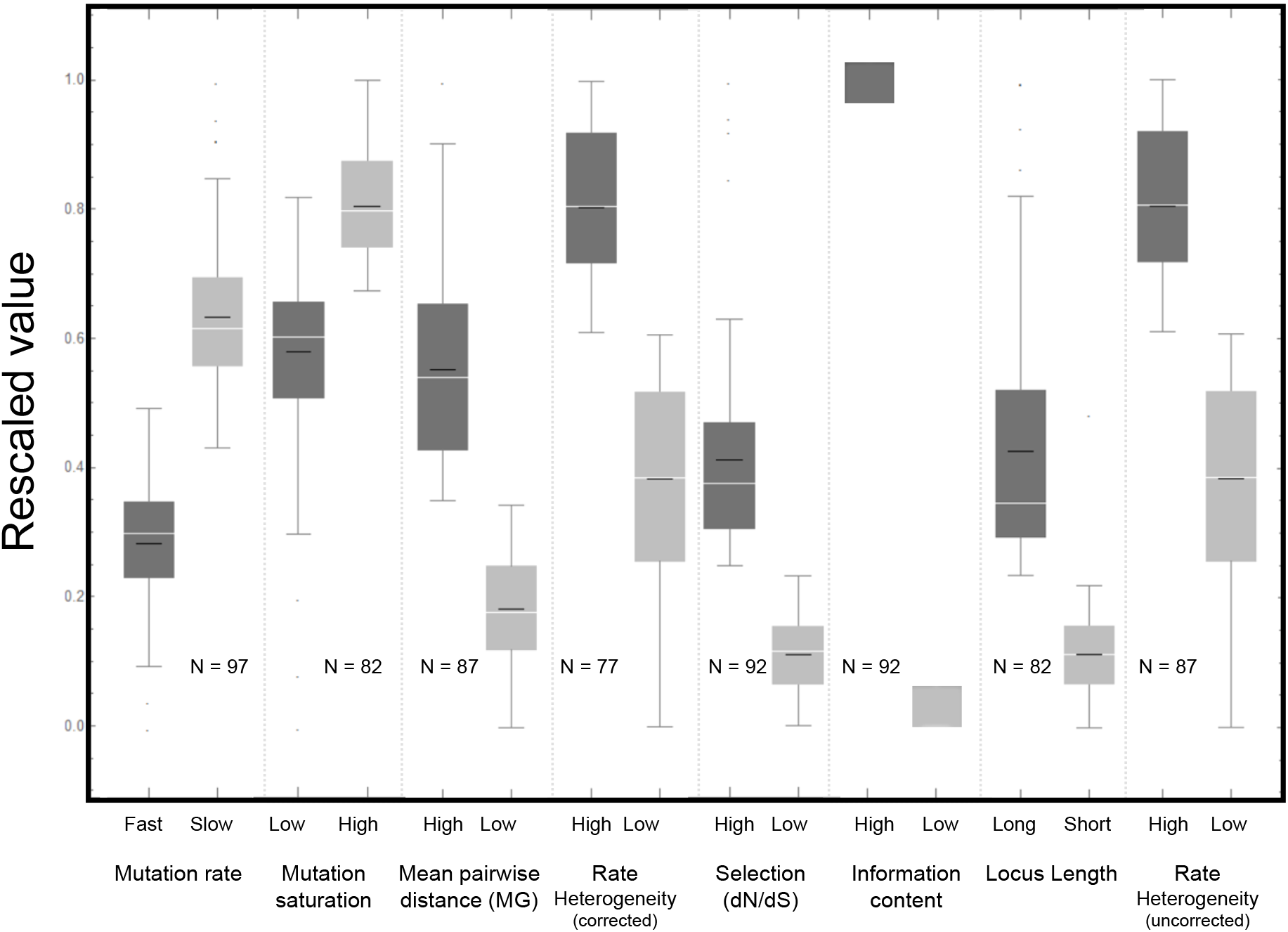
Distributions of feature values tested for phylogenetic utility. Box-plots show the distribution of values (rescaled between 0 and 1) for both treatments of eight factors tested: mutation rate, mutation saturation, mean pairwise sequence distance, among site rate heterogeneity corrected for locus length, selection (dN/dS), a prori information content, locus length, and total among site rate heterogeneity. Boxes represent middle 75% quantiles; whiskers represent the remaining quantiles with points being outliers. White lines are the median and black lines are the mean. N indicates how many loci are in each treatment.

### Experimental Design in RADICAL

To assess stability of performance to various scales of sampling, we utilized the random addition concatenation of loci (RADICAL) described in Narechania et al. (2012). This was implemented using the “radicalRun” function in Phyloinformatica v. 0.93. Within RADICAL the fast tree reconstruction method (options -fast, -ntop 10) in IQ-TREE (Nguyen et al., 2015) was utilized. To overcome the low data completeness for some taxa, the first step of this implementation chose the three alignments with the most taxa and concatenates them with nine randomly drawn alignments. Subsequently, five loci were added at random for each additional step until all loci are sampled. A starting BIONJ tree was used in the first step (option -t BIONJ) and subsequent steps use the final tree from the previous step as an initial tree. The GTR+G model was used in an unpartitioned analysis parallelized over two cores. This was repeated for ~100 iterations for each treatment.

To assess the optimality of the test subsamples we compared their performance in recovering the topologies inferred from all loci (265_Full, 265_2nd, 265_Reduced, 265_Full_IQ). This was measured as the mean of all Robinson-Foulds (RF) distances (Robinson and Foulds 1981) from the RADICAL tree from each step at each iteration to the four trees. We use a set of baseline trees, as opposed to a single baseline, to establish: (i) a range of RF distances that could be considered reasonably accurate and (ii) as an acknowledgement of the probable inaccuracy of any single baseline tree. We plotted the means and fit with exponential models, which Narechania et al. (2012) showed was the shape of typical RADICAL curves.

To assess precision of tree estimation, we plotted RF comparisons among all trees within a RADICAL step. We used a linear best-fit model for this data, as opposed to the exponential model, to accommodate the expectations that some RF values could equal 0.

We compared distributions of RF distances among the test subsampling treatments to assess the differences among subsamples. We compare each of the RADICAL tests at the 14^th^ step (the latest step that is not the last step for any treatment) and the last step, which showed identical patterns of statistical significance. All statistical comparisons were done with a Z-Test, which is not sensitive to slight deviations from normality with sample size greater than 40 (Ghasemi and Zahediasl 2012).

### Final Tree Searches

The results from RADICAL identified five features demonstrating high phylogenetic synecdoche and showing benefits to tree precision. We selected two sets of 100 loci using the factors shown to improve RADICAL curves (length-corrected rate heterogeneity, mean pairwise sequence distance, mutation rate, saturation, and selection). To do this we log-transformed and rescaled the values between zero and one, and altered them so that higher values corresponded to the more desirable feature (i.e. low rate heterogeneity, low pairwise sequence distance, low selection, low saturation and high mutation rate). These were then added together and treated as a score for each locus. For the first set of 100 loci, we simply concatenated the 100 highest scoring loci (65,798 nucleotides long; “100_Full” alignment). For the second set, we added the scores to a log-transformed and rescaled version of the number of taxa with data for a given locus (i.e. completeness). The 90 top scoring loci were chosen and concatenated along with the 10 next most complete loci (83,822 nucleotides long; “C100_Full” alignment). The new alignments had only 38% as many loci and ~1/3 as many nucleotides as the 265LociFull alignment.

Final tree searches and bootstrapping were performed on the 100_Full and C100_Full alignments in RAxML v. 8.2 using the same protocol, partitioning and node support strategy discussed above with 100 independent tree searches. Searches were again performed on three conformations of each alignment: 1) all 100 loci (“Full”), 2) only 2^nd^ codon positions (“2nd”), 3) only positions present for more than 50% of taxa (“Reduced”).

## Results

Our experimental comparison of six features (Fig. 2) using the random-addition concatenation (RADICAL) process (Fig. 3) showed that fast mutation rate, low saturation, low mean pairwise sequence distance, low rate heterogeneity, and low selection all contribute to faster convergence to the full data tree topologies (P-values all << .005; Fig. 4A; also see, Fig. S4.2). Loci with high rate heterogeneity, which tend to be longer, improved tree recovery but only when not corrected for locus length (Fig. S4.4). Fast mutation rate, low mean pairwise sequence distance, low rate heterogeneity, low selection, and low information content all result in more precise estimation of trees (P-values all << .005; Fig. 4B; also see, Fig. S4.3). Saturation had no statistically significant effect on tree precision.

**Figure 3.**
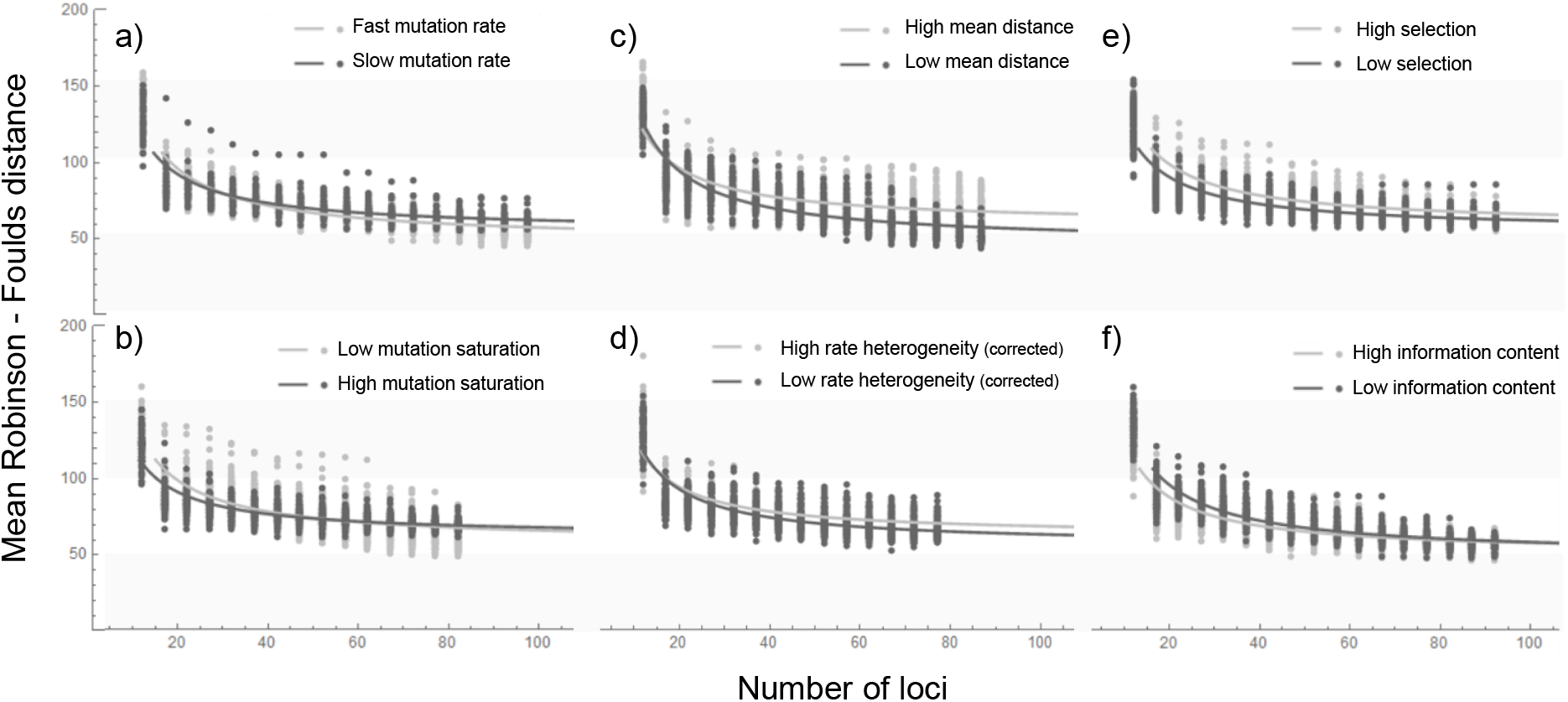
RADICAL curves for six tests of phylogenetic utility. Dots represent the mean Robinson-Foulds distance to four baseline trees. Lines show best fit exponential curves.

**Figure 4.**
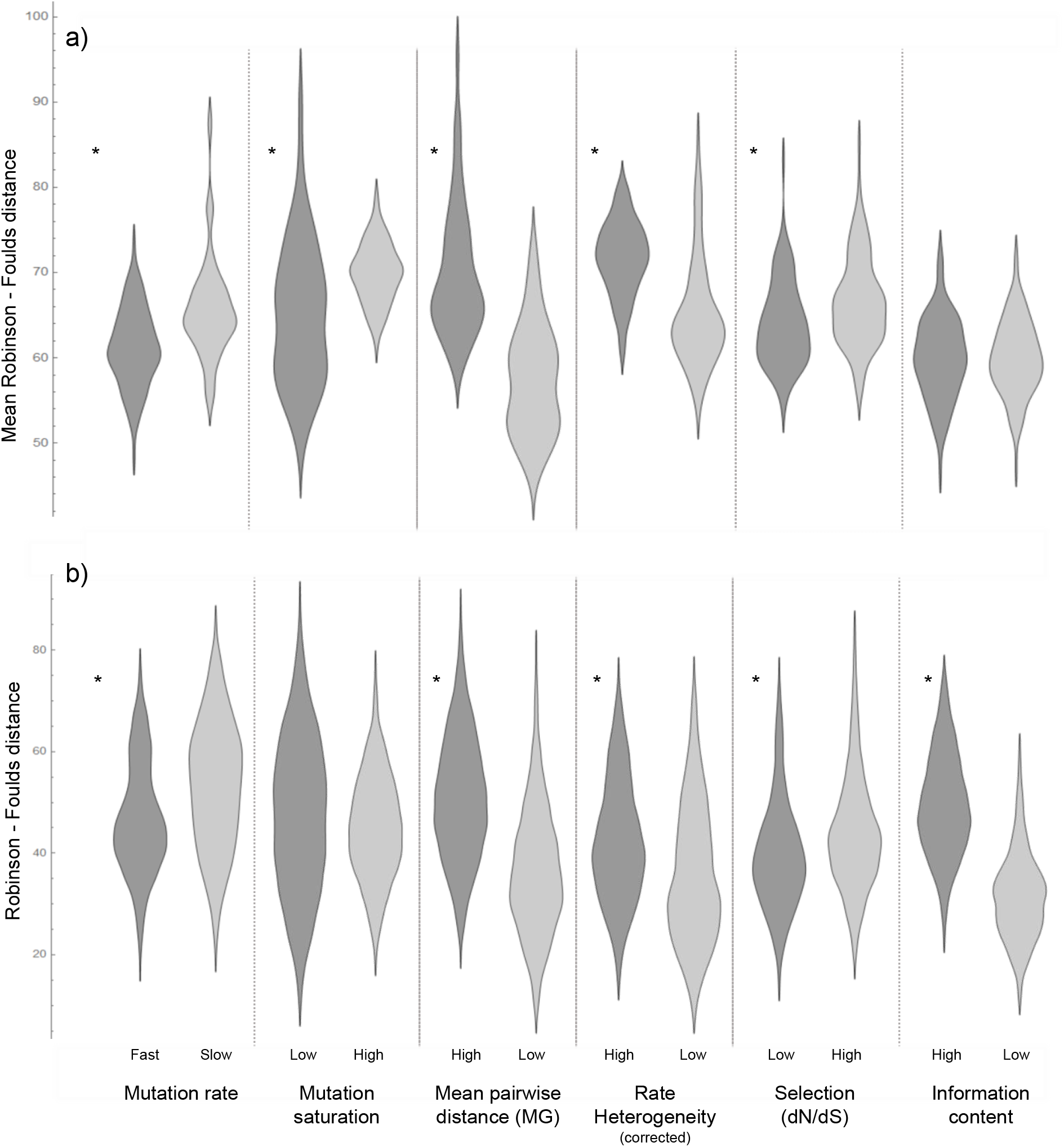
Distribution plots showing (A) mean similarity (RF distance) to baseline trees and (B) similarity among tree inferences of two treatments for six factors. Each comparison is from the 14^th^ concatenation step of each RADICAL run. Asterisks indicate a statistically significant difference, as determined by a Z-Test of mean comparisons (alpha < 0.005).

The two trees inferred from the optimized subsamples of 100 loci were 48 (100_Full) and 20 (C100_Full) RF units of distance away from the 265_Full tree (Table 1). For context, the 265_2nd and 265_Reduced trees were 60 and 56 RFs different, respectively. C100_Full tree had the lowest distance to 265_Full tree (Table S5.1) and is in the 99.95^th^ percentile of all the 23,740 comparisons to the baseline trees done in RADICAL (100_Full tree’s distance was in the 88^th^ percentile). The 100_Full and C100_Full trees also had slightly longer internal branch lengths compared to external branch lengths (i.e., decreased leafiness; Table 1). The mean bootstrap support and internode certainty of 100_Full and C100_Full were slightly lower than those in 265_Full (Table 1). The only trees deemed plausible given an alignment were 265_Full, 100_Full, C100_Full, and C100_Reduced (Table 1). The C100_Full phylogeny was the only tree deemed plausible considering more than one alignment.

**Table 1.** Comparison of tree quality and support among nine trees.

| Tree | Leafiness<sup>a</sup> | RF<sup>b</sup> | ln L<sup>c</sup> | Mean tree bootstrap support | Relative tree certainty | # of AU tests passed<sup>d</sup> |
| --- | --- | --- | --- | --- | --- | --- |
| 265_Full | 0.171 | 0 | -3038176.6 | 92.1 | 0.81 | 1 |
| 265_2nd | 0.214 | 60 | -433293.3 | 73.7 | 0.51 | 0 |
| 265_Reduced | 0.151 | 56 | -793673.2 | 87.4 | 0.75 | 0 |
| 100_Full | 0.154 | 48 | -919296.0 | 87.9 | 0.72 | 1 |
| 100_2nd | 0.179 | 98 | -117967.0 | 60.9 | 0.34 | 0 |
| 100_Reduced | 0.151 | 66 | -444162.8 | 85.9 | 0.70 | 0 |
| C100_Full | 0.164 | 20 | -1207800.6 | 89.7 | 0.74 | 2 |
| C100_2nd | 0.198 | 96 | -159517.2 | 65.4 | 0.41 | 0 |
| C100_Reduced | 0.157 | 44 | -501350.8 | 87.8 | 0.74 | 1 |

<sup>a</sup> The ratio of total tip branch lengths to internal branch lengths.<sup>b</sup> Robinson-Foulds distance to the 265\_Full tree<sup>c</sup> Log-likelihood of the tree given its alignment<sup>d</sup> how many times the tree was accepted ( $p > 0.05$ ) by an AU test (maximum of three).

The C100_Full and 100_Full subsamples are composed of loci with qualities shown to be more optimal under the criterion of phylogenetic synecdoche. Also, both 100 loci trees display a lower ratio of total external branch length to internal branch length compared to the 265_Full phylogeny − implying a higher proportion of characters are supporting the relationships (Table 1) (i.e., more signal; Dornburg et al. 2018).

Topological distance between the subsample trees and the full data tree is an approximation for topological improvement. The C100_Full and 100_Full trees had distances of 20 and 48 RFs to the 265_Full respectively. We favor the C100_Full tree (Fig. 5) because its accounts for data completeness (which is known to improve phylogenetic recovery; Dell’Ampio et al. 2014), it recovered the best ∆lnL in an approximately unbiased (AU) test of tree support (Table S5.3), and was the only tree accepted by more than one AU test (Table 1; also, see Table S5.2, S5.3).

**Figure 5.**
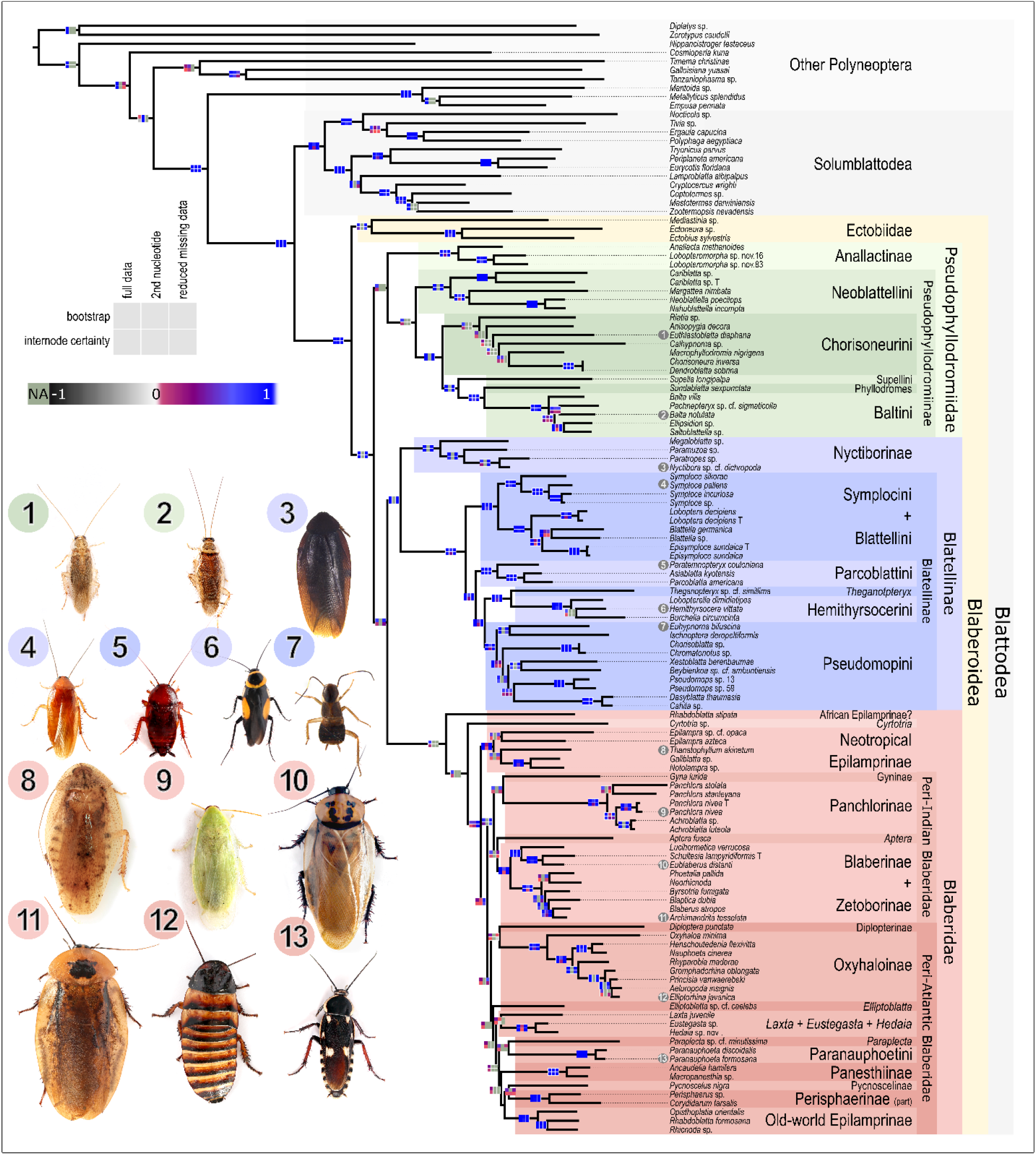
Phylogeny of Blaberoidea (C100_Full tree). Phylogeny of Blaberoidea, as inferred from a partitioned RAxML tree inference from 100 highest scoring and taxon complete loci. Support values in color-coded Navajo rugs are bootstrap frequency and internode certainty scores calculated from three trees, as described in the legend. “NA”/pink indicates the particular bipartition does not appear in the specified tree. Branch lengths are proportional to mutations. Major nodes are annotated and taxa are colored by clade. Numbers correspond to tip taxa depicted in photographs. Photographs by Dominic Evangelista.

The C100_Full phylogeny supports a number of key relationships in Blaberoidea. This tree shows Ectobiinae (Ectobiidae) sister to all other Blaberoidea; Anallactinae (nom. nov.) sister to Pseudophyllodromiinae (together Pseudphyllodromiidae nom. nov.); Blattellinae + Nyctiborinae (Blattellidae sensu nov.) sister to Blaberidae; monophyly of Blaberidae; and most Blaberidae falling into either the “Peri-Atlantic” or “Peri-Indian” monophyletic clades. There is strong node support throughout Blattellidae and in parts of Pseudophyllodromiidae (with the exception being Chorisoneurini; Figs. 5, 6). In Blaberidae there is strong support for recent relationships but most deep relationships lack strong node support (Figs. 5, 6). The relationship *Rhabdoblatta stipata* as sister to the remaining Blaberidae is strongly supported by our data, although node support suffers in the displayed tree due to the presence of one rogue taxon (*Cyrtotria* sp.).

**Figure 6.**
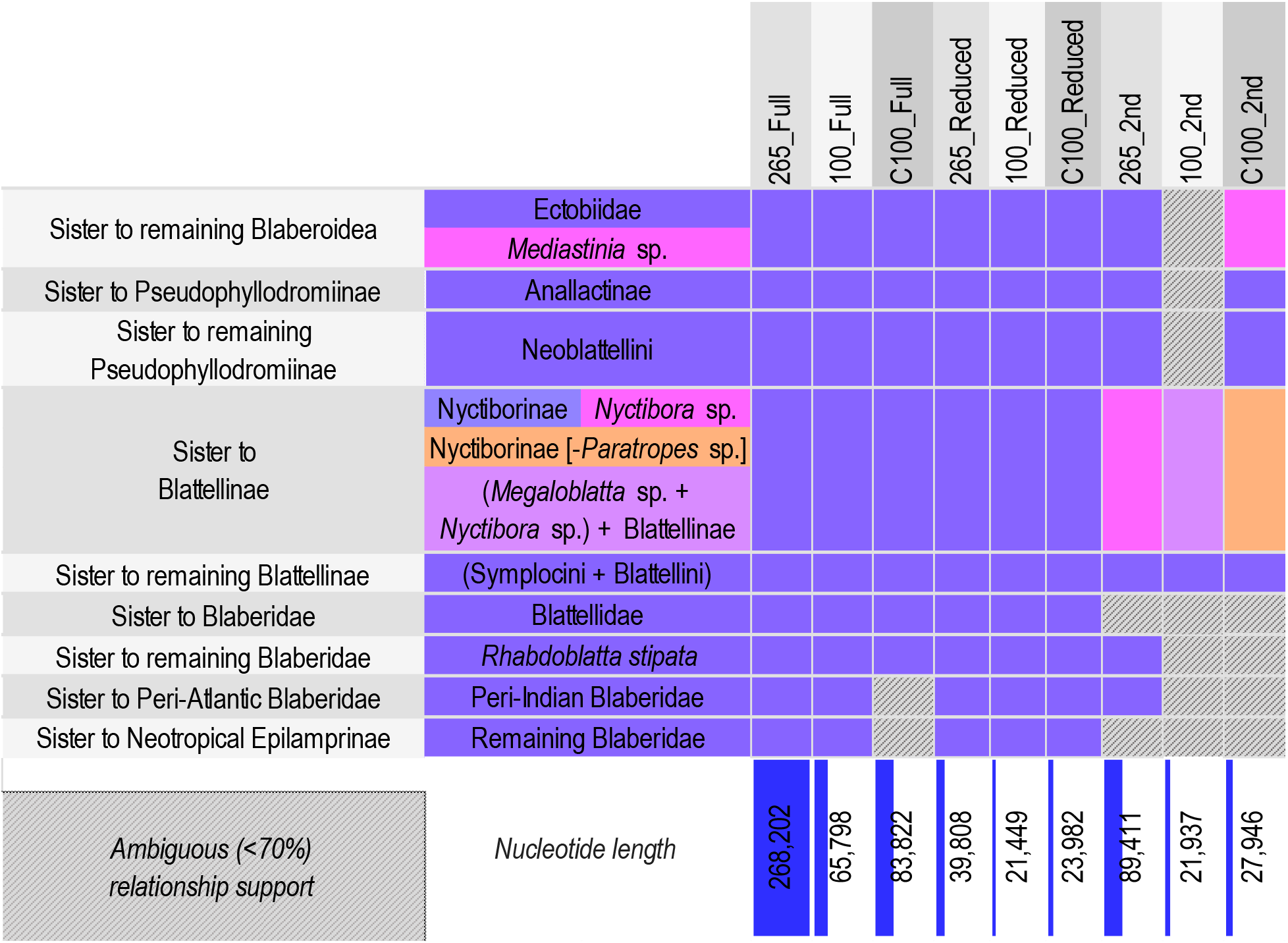
Heat map of support for selected backbone relationships in Blaberoidea. Support for relationships are given for the nine trees indicated in column labels. Alternative proposed relationships are indicated by cell coloration. Only relationships with more than 70% support are consider unambiguous and colored solidly [see Evangelista et al. (2018) for the method of calculating support for relationships from bipartition support values]. Nucleotide length of each alignment is shown in the bottom row, and the cell is colored proportional to the values.

## Discussion

*A priori* information content of loci was not a predictor of phylogenetic performance, which is surprising but not unprecedented (Chen et al. 2015). This is likely not a failure of the methods (i.e., MARE and SAMS) to infer information content but rather an inappropriate application of the methods. First, the subset of taxa we applied these methods to have been insufficient since taxon sampling impacts inferred character evolution (Venditti et al. 2006; Hugall and Lee 2007). Second, MARE intentionally scores a locus as being uninformative if there is a high proportion of missing data, and missing data patterns chang after new sequences are added. Third, single genes may not contain enough characters to inform phylogenetic information estimators. Assignment of character state changes as autapomorphies, synapomorphies or symplesiomorphies depends on the number of available characters. Finally, the methods could be confounded by long branch attraction (LBA), mistaking saturated loci for highly tree-like loci and wrongly attributing homoplasy to split support. Although our final tree does not show exceedingly long branches, all the reconstructed trees exhibited features of other long branch effects [abundant short internodes (Fig. 5 and Fig. S3.1) and a high leafiness (Table 1)] (Wägele and Mayer 2007; Kück et al. 2012).

While high information content loci performed poorly, fast evolving and low saturation loci converged rapidly towards the expected topology. This result contradicts the idea that saturation is a consequence of high evolutionary rate, two features that were not highly correlated (R = −0.11; Table S1.1, S1.2) in our dataset. This also contrasts with the assumption that slow evolutionary rates are assumed to be superior (Chen et al. 2015). Although the superiority of slow rates are only supported when fast evolutionary rate result in saturation and the optimal rate changes depending upon the phylogenies’ shape (Dornburg et al. 2018). Unsaturated but fast evolving loci should have a higher density of phylogenetic information than slow characters.

Low rate-heterogeneity among loci improved tree inference (Figs. 2, 3). This is consistent with studies showing that rate heterogeneity confounds the assumptions of phylogenetic models (Galtier et al. 2006; Frandsen et al. 2015) and better adherence to those models improves phylogenetic inference (Kjer and Honeycutt 2007; Doyle et al. 2015; Reddy et al. 2017). However, total high rate heterogeneity is not problematic (Fig. S4.4) since a higher variance in rates can be a result of loci being longer (Table S1.1).

Strong stabilizing selection (Figs. 2, 3), as defined by low mean ratio of non-synonymous to synonymous mutations (dN/dS), also improved tree reconstruction. This agrees with higher mean locus dN/dS corresponding to: more non-synonymous sites under positive selection, which are difficult to model (discussed in Beaulieu et al. 2018); and purifying selection of synonymous mutations (Spielman and Wilke 2015), which yield compositional bias and phylogenetic errors (Cox et al. 2014).

The best performing loci had low mean pairwise sequence distance (Figs. 2, 3). Sequence distance is correlated to a suite of other features also related to phylogenetic utility (Bai et al. 2013; Struck 2014; Borowiec et al. 2015; Chen et al. 2015; Lewis et al. 2016; Borowiec 2019). In particular, decreasing mean pairwise sequence distance may be similar to minimizing saturated loci because they both reduce the probability of LBA (Struck 2014). LBA can lead to rogue taxon placement, which our analysis is particularly sensitive to due to usage of the RF metric (Kuhner and Yamato 2015).

Our experimental design specifically attempts to control for extraneous features (see Figs. S1.1, S4.2, S4.4), something similar studies omitted. We paid particularly close attention to locus length which was moderately correlated with rate heterogeneity, mean pairwise sequence distance, and saturation (Table S1.2). Yet, these treatments with longer loci did not perform better (i.e., Fig. S4.4). Thus, more effectively controlling for locus length may yield a stronger effect-size.

### Phylogeny of Blaberoidea from an Optimized Subsample

We found strong support for a number of key relationships in Blaberoidea (Fig. 6). Note that support values from the 100_2ndC100_2nd and 100_ReducedC100_Reduced alignments in Figure 5 are misleading due to *Cyrtotria* sp., whose placement became highly volatile after further data reduction. Our discussion here consider support for relationships with this species removed (see supplemental data).

The C100_Full phylogeny of Blaberoidea (Fig. 5) largely agreed with the phylotranscriptomic study of Evangelista et al. (2019), and the taxa shared among them have identical relationships but with lower support values here. We found strong support for Ectobiinae (constituting *Ectobius sylvestris, Mediastinia* sp. and *Ectoneura* sp.) as the sister-group to the remaining Blaberoidea, which was also recovered in Wang et al. (2017). This was not recovered in other recent studies (Djernæs et al. 2015; Legendre et al. 2015) but Evangelista et al. (2018) showed that it was still plausible. We recovered a clade containing *Anallacta methanoides* as sister to Pseudophyllodromiinae. Evangelista et al. (2019) discussed molecular and morphological support for this novel grouping. Pseudophyllodromiinae + Anallactinae was recovered as sister to the remaining Blaberoidea with the exception of Ectobiinae, which is supported by phylotranscriptomics (Evangelista et al. 2019) but is not seen in most other studies. Blattellinae was sister to Nyctiborinae (as in Klass and Meier 2006; Djernæs et al. 2015; Legendre et al. 2015; Evangelista et al. 2019), which in turn was sister to Blaberidae − a hypothesis supported by morphology (Klass and Meier 2006) and molecular data (Bourguignon et al. 2018; Evangelista et al. 2019). The recovered phylogeny of Blaberidae is largely a unique hypothesis compared to the most recent works done (Legendre et al. 2015; Legendre et al. 2017; Bourguignon et al. 2018) but all the studies, including ours, have low support for deep relationships. For the first time, and with moderately strong support, we recovered an African *Rhabdoblatta* as the sister-group to the remaining Blaberidae. Also with strong support, Neotropical Epilamprinae was sister to the remaining Blaberidae with the exception of African *Rhabdoblatta*. We recovered a Neotropical and African clade (Gyninae + Panchlorinae + *Aptera* + Blaberinae + Zetoborinae) with moderate support, as in Bourguignon et al. (2018). This was sister to an old-world clade (Diplopterinae + Panesthiinae + Perisphaeriinae + Oxyhaloinae + old-world Epilamprinae + Pycnoscelinae + Paranauphoetinae), but the latter clade, which has never been recovered in any previous phylogenetic study, had very low support throughout. Further discussion of recovered phylogenetic relationships is given in the Supplementary Material Section 4.

While support for the specific relationships listed above are comparable among the 265_Full and C100_Full trees, there are strong differences among these topologies (Fig. S5.1). Unfortunately, most of these topological differences cannot be interpreted in light of previous morphological or molecular studies because comparative data is lacking for these clades. One separate study (Evangelista et al. 2019 unpublished data) is examining evidence among the more volatile relationships of Blaberidae so we refrain from addressing those here.

While this is the first time many of these relationships have been recovered with strong signal a number of improvements need to be made in the future. Increased taxon sampling and further optimization of the dataset could further improve tree quality, particularly for recent relationships in Blaberidae. Finally, applying more robust tests of topological support such as four-cluster likelihood mapping (Strimmer and Haeseler 1997), other quartet methods (Pease et al. 2018) and better handling rogue taxa (Aberer et al. 2013) could further improve our identification of robust relationships and dark nodes.

### Practical Improvements to Genomic Systematics

Reducing dataset size has practical implications for phylogenetic projects. Computation time for alignment estimation and topology testing (AU test) is reduced proportionally to the reduction in dataset size (Table 2). Gains in efficiency of phylogenetic inferences are 4-6 fold improvements (Table 2). These are significant in practice. For instance, the C100_Full tree only took 275 CPU hours to reach the optimization stop criteria, over a month faster than 265_Full (1025 CPU hours). Reducing heterogeneous loci provides the ability to simplify the evolutionary mode − avoiding over-parameterization (Sullivan and Joyce 2005) and further improving computation time. We did not re-estimate optimal partition and model schemes for our dataset so we could consistently compare our results with the 265_Full phylogeny. Yet, doing so would be a qualitative improvement (e.g., Philippe and Roure 2011).

**Table 2.**
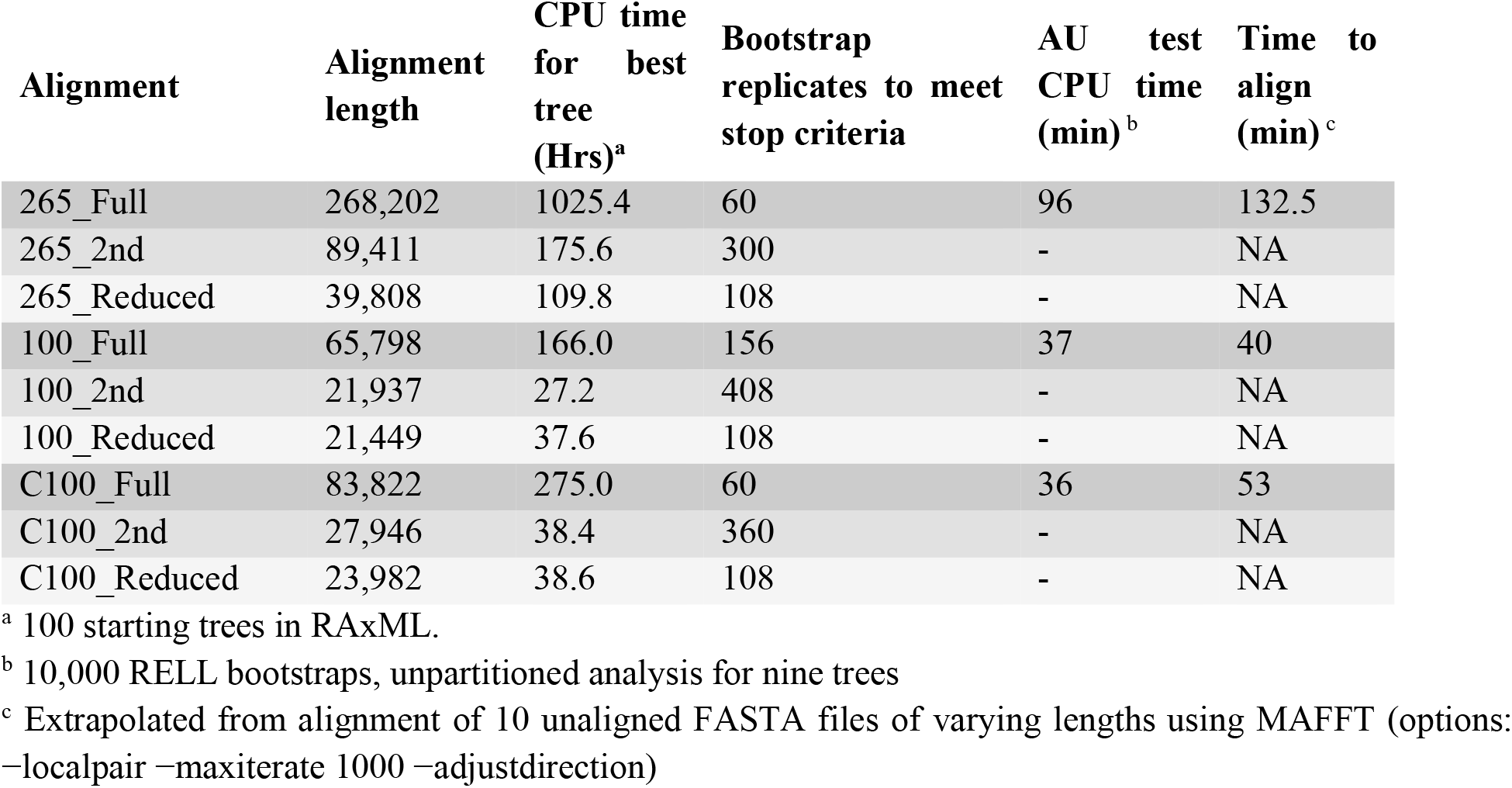
Comparison of various activities derived from three alignment conformations of 265 loci and two sets of 100 loci.).

Reducing computational effort of tree inference is extremely important considering that phylogenetic studies rarely do such analyses only once. For instance, we conducted two preliminary tree inferences and two quality checking inferences, each of which involved analyzing the 265_Full alignment. When adding together main tree searches, those of different alignment conformations and assessing support through resampling methods, computation time is a limiting resource even when having access to multiple world-class supercomputing clusters (e.g., Shen et al. 2017 and this study). Improving the practicality of inferring trees and their support makes the application of downstream analyses (e.g., divergence date inference, diversification analyses, inferring trait evolution) more feasible as well. The important caveat, though, is one must have a large preliminary dataset from which to select the most optimal loci. We provide a user-friendly summary of our recommended approach in Supplementary Material S.6.

### Conclusions

Here we used the criterion of phylogenetic synecdoche to identify optimal subsamples which in turn improved phylogenetic inference. High evolutionary rate, low saturation, low mean pairwise sequence distance, low rate heterogeneity, and low selection were the most optimal traits of loci. Measures of *a priori* phylogenetic information content that rely on split signal or tree-likeness may not be meaningful for short alignments or limited taxon samples. These findings are consistent with past studies and phylogenetic theory (Townsend 2007; Cox et al. 2014; Doyle et al. 2015; Dornburg et al. 2018), and provide the opportunity to meaningfully target maximally informative loci using locus capture methods (Lemmon et al. 2012; Brandley et al. 2015; Gilbert et al. 2015). Subsamples reduced by two thirds of total data length and optimized under the above specifications exhibited recovery of a phylogeny exceptionally similar to the tree inferred from all the data, with topological improvements approximated at 20 Robinson-Foulds units on our tree of 124 tips. This demonstrates that by targeting maximally phylogenetically useful loci, one can drastically reduce the monetary cost and computational resources of projects while maintaining, or even improving, the quality of the results. Thus, in addition to robustly recovering the phylogeny of Blaberoidea, this study provides an efficient workflow to infer robust phylogenomic trees with optimized datasets. The only potential drawback is decreased node support and lessened ability to do further data reduction tests.

## Availability of data and material

Transcriptome datasets were taken from Evangelista et al. (2019) and newly sequenced data is deposited on the NCBI Sequence Read Archive (SRP155429). Other analyzed files are available on the Dryad digital repository (doi:10.5061/dryad.9mf1pr7).

## Authors’ contributions

DAE, FL and AYK obtained funding for the study. DAE and FL conceived the study and organized the taxon sampling design. DAE ran preliminary analyses and designed genomic sampling with assistance from SSi and BW. DAE, MMW and JLW designed the molecular baits. DAE extracted and enriched genomic DNA, with MMW, JLW, and MKK providing guidance. DAE did all bioinformatics, wrote custom software, executed all analyses, with guidance from FL. DAE, OB and FL revised taxonomic descriptions of taxa with assistance from BW. DAE and FL wrote the paper with assistance from SSi, AYK and BW. DAE and BW composed the figures. SSi, BW, JLW, MKK, and OB, provided early access to transcriptome datasets and contributed in analyses and curation of those datasets as well as feedback to the manuscript.

## Acknowledgements

Thanks to the 1KITE consortium who supported this research with preliminary data and advice with software. Specifically, appreciation extends to Karen Meusemann, Alexander Donath, Bernhard Misof, Xin Zhou, Shanlin Liu, Ralph S. Peters, Lars Podsiadlowski, Ward Tollenaar, Mari Fujita, and Ryuichiro Machida. Huge thanks to all breeders (Nicolas Rousseaux, Tristan Shanahan, T.J. Ombrelle and Piotr Sterna), colleagues (Mike Picker), museums (MNHN, MFN, NHMUK and CAS) and curators (Jurgen Deckert, and George Beccaloni) who assisted in providing specimens. Great appreciation to New England Biolabs, MycroArray (now Arbor Biosciences), Sara Ruane, Ciara-Mae Mendoza, Melissa Sanchez-Herrera, Steven Ramirez and Mihaela Glamoclija for providing assistance in the lab. Additional thanks to Brian O’Meara for guidance and advice. This research could not have been completed without the support of NSF (award # 1608559), all other funding agencies, the MNHN - Paris, Rutgers University and the University of Tennessee - Knoxville.

