## Supplementary material for "Phylogenetic Synecdoche Demonstrates Optimality of Subsampling and Improves Recovery of the Blaberoidea Phylogeny" (from another version of this preprint: version 1, 2019-04-07)

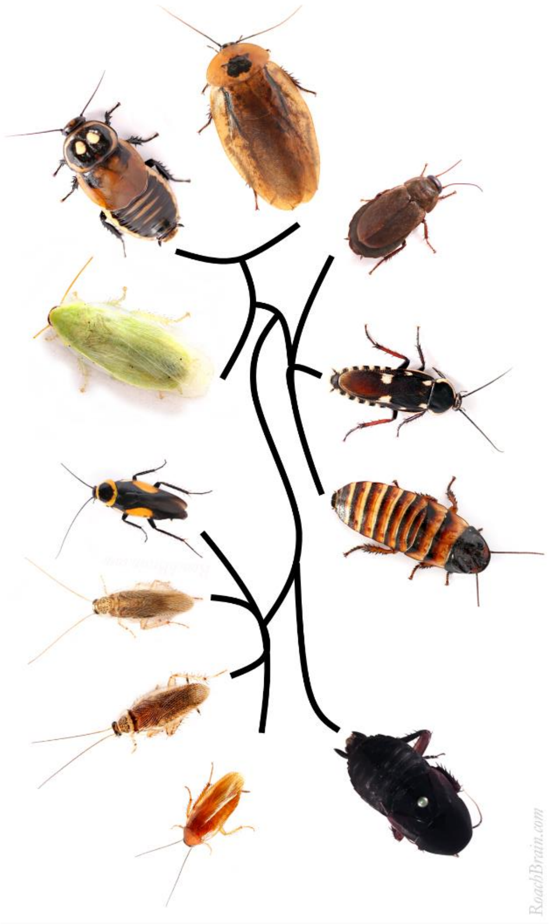

Supplement
to

### Phylogenetic synecdoche demonstrates optimality of subsampling and improves recovery of the Blaberoidea phylogeny

*Published in BioRxiv by*

**Authors**

Dominic A. Evangelista, Sabrina Simon, Megan M. Wilson, Manpreet K. Kohli, Jessica L. Ware, Akito Y. Kawahara, Benjamin Wipfler, Olivier Béthoux, Philippe Grandcolas, & Frédéric Legendre

5 - April - 2019

### Section 1: Supplemental methods

#### Correlations among locus traits

Many traits of the calculated traits among the 265 loci were strongly (Spearman's coefficient of correlation, "R", >= 0.5), moderately (R between 0.5 and 0.35) or weakly (R between 0.35 and 0.2) correlated with one another (Table S1.1). In particular, rate heterogeneity and alignment length are strongly correlated (even after correcting for nucleotide length of each locus). Corrected rate heterogeneity, mean pairwise sequence distance and mutation saturation correlate with all traits considered except for information content, evolutionary rate, and selection. Information content and mutation rate, contrastingly, did not correlate with any other trait except (weakly) with number of taxa in the alignment. Finally, selection (dN/dS) was only correlated (weakly) with number of taxa in the alignment and nucleotide compositional bias (RCFV). The abundant correlation among traits highlights the necessity for designing controlled experiments.

##### Table S1.1

*Symmetrical matrix of correlation coefficients among ten traits of 265 loci considered in this study. This symmetrical matrix shows the coefficient of correlation (Spearman's R) among ten traits for all loci. The cells are shaded blue with increasing values of correlation.*

| | Mean rate | Saturation | Mean pairwise seq. distance | # of rate categories/ length | Selection (dN/dS) | Information content | Nucleotide length | Total # of rate categories | # of total taxa | Total RCFV |
|---|---|---|---|---|---|---|---|---|---|---|
| Mean rate | 1.00 | -0.11 | -0.03 | 0.05 | -0.11 | 0.14 | -0.06 | -0.11 | -0.36 | -0.05 |
| Saturation | -0.11 | 1.00 | 0.72 | -0.64 | 0.04 | -0.09 | 0.64 | 0.59 | 0.42 | 0.48 |
| Mean pairwise seq. distance | -0.03 | 0.72 | 1.00 | -0.43 | -0.02 | 0.03 | 0.58 | 0.56 | 0.30 | 0.55 |
| # of rate categories/ length | 0.05 | -0.64 | -0.43 | 1.00 | 0.16 | 0.01 | -0.50 | -0.27 | -0.24 | -0.20 |
| Selection (dN/dS) | -0.11 | 0.04 | -0.02 | 0.16 | 1.00 | 0.09 | 0.04 | 0.04 | -0.23 | 0.31 |
| Information content | 0.14 | -0.09 | 0.03 | 0.01 | 0.09 | 1.00 | 0.11 | 0.07 | -0.21 | 0.07 |
| Nucleotide length | -0.06 | 0.64 | 0.58 | -0.50 | 0.04 | 0.11 | 1.00 | 0.94 | 0.22 | 0.25 |
| Total # of rate categories | -0.11 | 0.59 | 0.56 | -0.27 | 0.04 | 0.07 | 0.94 | 1.00 | 0.25 | 0.24 |
| # of total taxa | -0.36 | 0.42 | 0.30 | -0.24 | -0.23 | -0.21 | 0.22 | 0.25 | 1.00 | 0.12 |
| Total RCFV | -0.05 | 0.48 | 0.55 | -0.20 | 0.31 | 0.07 | 0.25 | 0.24 | 0.12 | 1.00 |

#### Controlling for trait correlation

We took extensive lengths to control for correlation among traits in our RADICAL analysis by carefully designing treatment datasets. Each treatment's mean trait value and coefficient of correlation for extraneous traits are shown in Table 1. Figure S1.1 shows a principle components analysis for easy visualization of trait distributions. The overlap of the point clouds indicate that distributions of extraneous traits are largely equal among treatments.

##### Figure S1.1

*Principle components analysis (PCA) of locus traits in all treatments. Traits shown are: (A) mutation rate, (B) mutation saturation, (C) mean pairwise sequence distance, D) length corrected rate heterogeneity, E) selection (dN/dS), and F) information content. In the PCA for each treatment, the treated trait is omitted. Each point represents one locus. First two axes of the principle components are plotted against one another.*

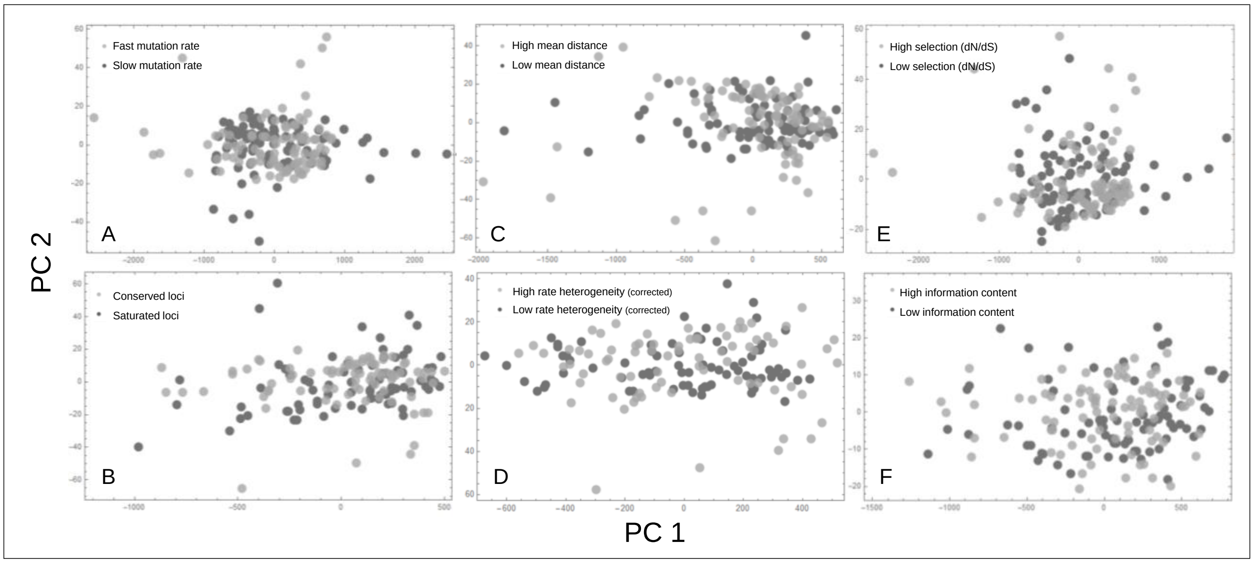

### Section 2: Taxonomic changes

#### New names for higher taxa

Our phylogeny and that of Evangelista et al. (2019) agree strongly that species of the genus *Anallacta* are more closely related to Pseudophyllodromiinae than to Blattellinae, despite contrary morphological evidence (Grandcolas 1996). We have found strong molecular support for a clade containing *Anallacta* and *Lobopteromorpha*. Given the morphological similarities of the two, we propose a novel subfamilial grouping.

Subfamily: Anallactinae nom. nov. Evangelista, Wipfler, and Legendre
*Systematic scope: Anallacta* Shelford, 1908, *Lobopteromorpha* Chopard, 1952, *Dipteretrum* Rehn, 1922

*Differential diagnosis*: According to, members of this subfamily can be distinguished from all other Blaberoidea by the synapomorphies: (i) left paratergite of sternite 8 with an elongated process (as opposed to without a process); (ii) hooked phallomere of male genitalia on the left side (as opposed to on the right). This character can be used in combination with the following plesiomorphy to delimit the clade: (iii) male genital sclerite l3d *sensu* Grandcolas (1996) not ring shaped (as opposed to ring shaped).

*Remarks*: Character (i) may or may not be articulated in Anallactinae. Bohn (2007) discusses a similar process as character (i) found in *Supella* and *Metabelina*, although it is mentioned that their homology is unlikely. Evangelista et al. (2019) discuss characters (ii) and (iii) in their supplementary material with respect to *Anallacta* and its placement relative to Pseudophyllodromiinae and Blattellinae.

Other less well-defined or homoplasious characters that could be distinctive of the subfamily are: body sclerotization heavy (as opposed to light); typically dark in color (as opposed to buffy); tegmina brachypterous or just meeting the end of abdomen (as opposed to apterous or tegmina extending beyond the end of abdomen); terga 7 visibly modified (as opposed to not modified or modified on other segments; Roth 1969).

While Princis (1955) and Bonfils (1969) compared *Euloboptera* Princis, 1955 to *Lobopteromorpha* and other similar genera, *Euloboptera* are probably not Anallactinae. *Euloboptera* is missing the left process (character i), and there is no visible tergal modification (Princis 1963; Bonfils 1969). The same applies to *Neolobopteromorpha* Bonfils, 1969. We have not examined any specimens of these genera other than *Lobopteromorpha*. *Dipteretrum* is included in this group on the basis of *Dipteretrum rudebeckae* Princis, 1963, which possesses character state (i) (Princis 1963).

Our phylogeny (Fig. 5) and previous studies (Evangelista et al. 2019) show that *Loboptera* Brunner von Wattenwyl, 1865 and *Lobopterella* Princis, 1957 are deeply nested within Blattellinae. They also do not bear the relevant combination of diagnostic characters.

Unranked: Orkrasomeria nom. nov. Evangelista, Béthoux, Wipfler, and Legendre
Systematic scope: Blattellinae Karney, 1908, Nyctiborinae Saussure and Zehntner, 1893, and Blaberidae Saussure, 1864.

*Differential diagnosis*: According to Klass and Meier (2006), members of this clade can be distinguished from all other Blaberoidea by the following synapomorphies: (i) ootheca rotated inside vestibulum so keel is directed laterally (as opposed to not rotated so keel is directed dorsally); (ii) in male genitalia, sclerite L1 absent (as opposed to present); (iii) tendon tve present and distinct (as opposed to absent or vestigial); (iv) muscle l3 absent (as opposed to present); (v) muscle l36 present (as opposed to absent); (vi) muscle s14 present (as opposed to absent). Morphological terminology here follows Klass and Meier (2006). See that paper, Klass (1997) and Klass (2001) for descriptions of phallomeres and musculature.

*Etymology:* A modification of the greek translation for "extreme limits of bodies" − "ακραία όρια των σωμάτων". This refers to the extreme variation in body size of the clade. The clade contains the cockroaches with the largest wing-span (*Megaloblatta*), greatest body mass (*Macropanesthia*) and those with the smallest body size (*Attaphila;* Bell et al. 2007; Djernæs 2018). There are various other examples of divergent changes in body size throughout the clade (e.g., in *Ischnoptera*, Blaberinae+Zetoborinae, *Epilampra*, Oxyhaloinae).

#### New family combinations

Ectobiidae *sensu* Beccaloni and Eggleton (2011) has been hypothesized as paraphyletic since McKittrick (1964). Recent studies have all supported this hypothesis (Grandcolas 1996; Djernæs et al. 2012; Legendre et al. 2015; Wang et al. 2017; Evangelista et al. 2018; Evangelista et al. 2019). Grandcolas (1996) noticed this early on and proposed changes to compensate. We follow those changes here with some modification.

Family: Pseudophyllodromiidae Grandcolas, 1996 *sensu nov.*
Systematic scope: Pseudophyllodromiinae Vickery and Kevan, 1983 and Anallactinae subfamily *nov*.

*Differential diagnosis*: No morphological or behavioral synapomorphies or other characters are known to distinguish this group from other Blattodea.

*Remarks:* Grandcolas (1996) proposed this name to include Pseudophyllodromiinae Vickery and Kevan, 1983 but did not include *Anallacta* or any clade resembling Anallactinae subfamily *nov.* Therefore, the diagnostic characters relevant to Pseudophyllodromiidae sensu Grandcolas, 1996 alone are not relevant to the new scope of this taxon. See the diagnostic characters for both Anallactinae above, and those for Pseudophyllodromiinae in other publications (e.g., McKittrick 1964; Grandcolas 1996; Klass and Meier 2006) for morphological diagnosis of this taxon.

Family: Blattellidae Karny, 1908 *sensu nov.*
Systematic scope: Blattellinae Karney, 1908 and Nyctiborinae Saussure and Zehntner, 1893.

*Differential diagnosis*: According to Grandcolas (1996), members of this family can be distinguished from all other Blaberoidea by the following combination of putative synapomorphic character states: (i) in male genitalia hooked phallomere on left (as opposed to right); (ii) in male genitalia, sclerite R3d *sensu* Grandcolas (1996) elongated longitudinally (as opposed to transversely); (iii) in male genitalia sclerite R2 *sensu* Grandcolas (1996), cleft directed forward (as opposed to backward; see Grandcolas 1996); (iv) ootheca rotated inside vestibulum so keel is directed laterally (as opposed to not rotated so keel is directed dorsally). Note that character (i) also occurs in some Ectobiinae Brunner von Wattenwyl, 1865 (Bohn 1987).

*Remarks*: Beccaloni and Eggleton (2013) proposed that Ectobiidae Brunner von Wattenwyl, 1865 took precedence over Blattellidae Karny, 1908. Before this, Blattellidae was usually used to refer to Blattellinae, Ectobiinae, Nyctiborinae, and Pseudophyllodromiinae (syn. Plectopterinae), which has long been considered a paraphyletic group. Anaplectinae Walker, 1868 has also sometimes been considered as Blattellidae (or Ectobiidae) but this is a rogue taxon with no settled systematic position (Evangelista et al. 2018).

Family: Ectobiidae Brunner von Wattenwyl, 1865 *sensu nov.*
Systematic scope: Ectobiinae Brunner von Wattenwyl, 1865

*Differential diagnosis*: According to Roth (2003), the following character states are synapomorphies for Ectobiinae Brunner von Wattenwyl, 1865: (i) in subgenital plate, apodemes greatly elongated (as opposed to moderately elongated or lacking); (ii) on dorsal abdomen, 7$^{th}$ tergite modified into a fossa with setae (as opposed to other types of modifications, modifications of other segments, or lacking modification); (iii) subgenital plate with one or fewer styles (as opposed to two or more styles); (iv) in male genitalia, hooked phallomere highly elongated (as opposed to moderately long or stout). To these characters, we add the following synapomorphies: (v) in the hindwing plicatum *sensu* Brannoch et al. (2017), cross veins occurring between AA veins in a graded manner (as opposed to lacking cross veins, or cross veins un-graded); (vi) in hindwing plicatum, first AA vein with a single furcate anterior ramus that forms the boarder of the apical triangle and curves back posteriorly in apical most end, usually merged apically with the pseudostem (as opposed to having more rami, or rami not forming the border of the apical triangle, when present); and (vii) in forewing, CuA bent anteriorly (i.e. with a hump) in region just distal to arculus (as opposed to straight or slightly curved posteriorly). Note that wing branching terminology is based on Li et al. (2018) and wing vein homologies are as in Cui et al. (2018).

*Remarks:* We determined wing venation synapomorphies based on observation of European *Ectobius* sp. (IWCOB103) and published illustrations of Australian *Choristima* spp. (Roth 1992), *Ectoneura* spp. (Hebard 1943), and *Stenectoneura* spp. (Hebard 1943). We compared these against specimens of *Laxta* sp. (IWOOB703), *Periplaneta* sp. (IWCOB257), *Anaplecta* sp. (IWCOB1097), *Riatia* sp. (DEIWO002), and *Balta* sp. (DERU321). We also utilized published illustration of *Prosoplecta* sp. (Anisyutkin 2013) and original photos of specimens illustrated in Evangelista et al. (2015) (*Nyctibora dichropoda*) and Evangelista et al. (2019) (*Epilampra* spp., *Ischnoptera miuda*, *Euhypnorna bifuscina*, and *Buboblatta vlasaki*.

While Anaplectinae is currently hypothesized to be a Solumblattodea, its placement is uncertain (Djernæs et al. 2015; Evangelista et al. 2018). Thus, other positions of Anaplectinae could affect the polarity of some of the above character states. Specifically, character state (v) and (vii) are also found in *Anaplecta* sp. Also, the rami of the first AA vein may merge to form a boarder to the apical triangle (character vi, in part) in some species where the apical triangle forms an appendicular field (e.g., Anaplectinae, and some Pseudophylloromiinae). However, these species usually have more than two rami to the vein.

Princis (1971) used Ectobiidae to refer to Ectobiinae and Theganopteryginae. Grandcolas (1996) did not use the name Theganopteryginae but he agreed with Princis (1971) and included the “Theganopteryginae” genera in the subfamily Ectobiinae. Our phylogenetic topology recovers “Theganopteryginae” (*Theganopteryx*, *Burchelia*, *Hemithyrsocera*) deeply nested within Blattellinae. This is in following with the classification of Roth (2003). Thus, we consider Ectobiidae to be equivalent to Ectobiinae alone.

### Section 3: Supplemental discussion of the phylogeny of Blaberoidea

#### 265_Full tree

The tree we recovered from a RAxML analysis of the full alignment of 265 loci was relatively highly supported by bootstrap values. While we have proposed that the C100_Full phylogeny (shown in main text) is a more plausible phylogeny, we show the 265_Full phylogeny here for comparison (Fig. S3.1). The relationships among the two trees are the same without noteworthy differences.

#### Figure S3.1

*Phylogeny of Blaberoidea as inferred from a full alignment of 265 loci. Support values are illustrated as Navajo rugs and represent bootstrap frequencies and internode certainty scores from four analyses. The four analyses are: RAxML analysis of full 265 loci alignment, RAxML analysis of 2nd codons from 265 loci, RAxML analysis of 265 loci reduced to only the most complete positions, and IQ-TREE analysis of full 265 loci alignment. “NA”/Purple squares indicate that the displayed relationship did not appear in the particular tree. Major and minor taxonomic groupings are indicated.*

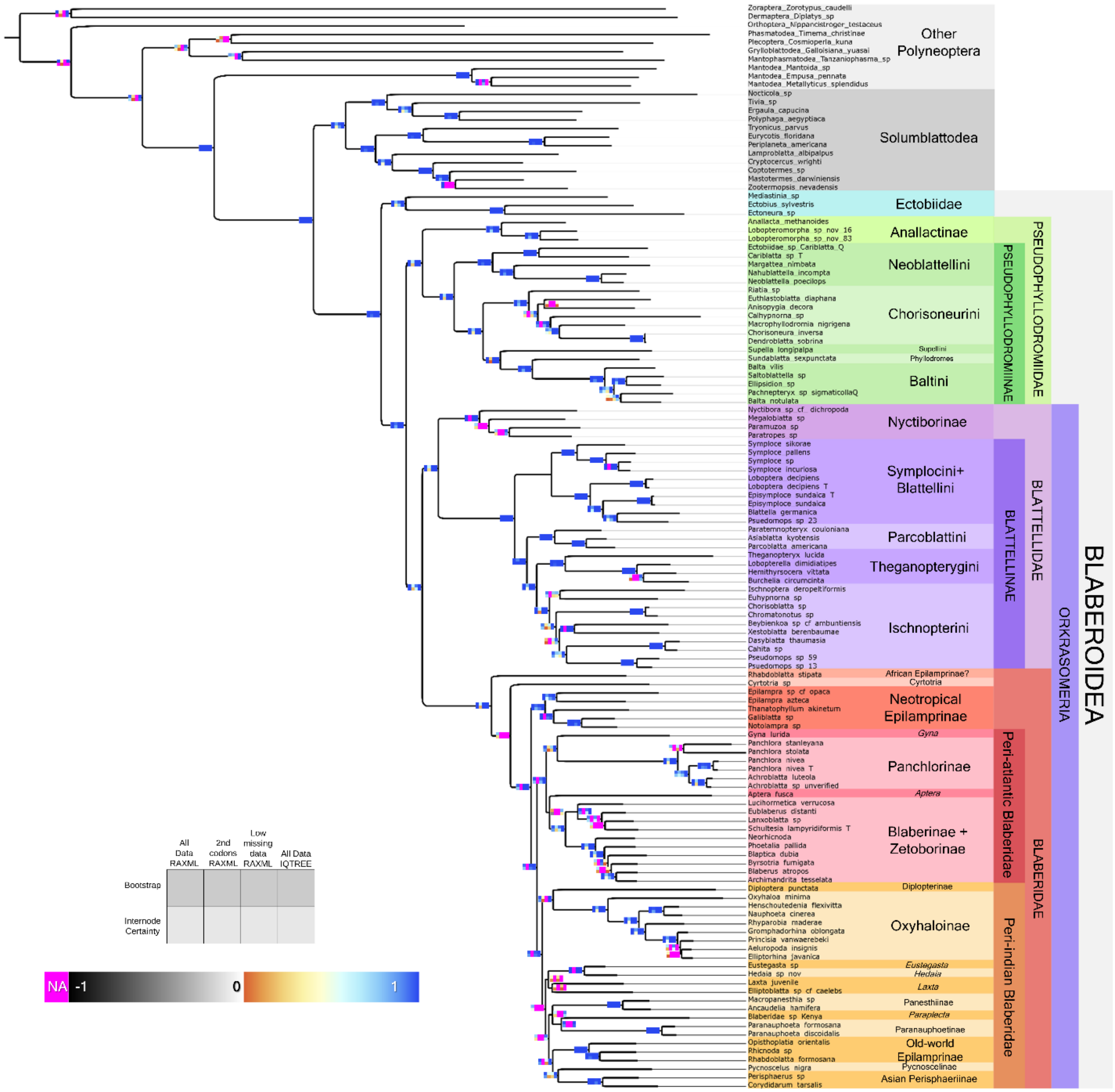

#### C100_Full phylogeny

The major insights into the phylogeny of Blaberoidea discerned from the C100_Full phylogeny are discussed in the main text, but we reserve a detailed discussion for here.

First, we recover Ectobiinae as consisting of *Ectobius*, *Ectoneura* and *Mediastinia*. *Ectoneura* was indeed recovered in Ectobiinae by Bourguignon et al. (2018), as expected based on its morphology-based classification (Bohn et al. 2010). However, *Mediastinia* was classified as Pseudophyllodromiinae (Hebard 1943), but this taxon is morphologically poorly defined.

We recover *Anallacta* in a clade with *Lobopteromorpha*. We recover this with strong support as the sister to Pseudophyllodromiinae, but both genera were previously classified as Blattellinae. Evangelista et al. (2019) also found *Anallacta* as sister to Pseudophylloromiinae and discussed morphological evidence for and against its placement in Blattellinae. We also find morphological evidence supporting this clade's monophyly as separate from Pseudophyllodromiinae and Blattellinae (Section 2). Thus, we elevate this clade to the subfamily level, calling it the Anallactinae (see Section 2). Together, the monophyletic clade comprising Anallactinae and Pseudophyllodromiinae we refer to as the family "Pseudophyllodromiidae" (see Section 2).

In Pseudophyllodromiinae, we recover three principle clades. Neoblattellini is recovered as sister to the remaining Pseudophyllodromiinae, with strong support. This clade constitutes mostly Neotropical taxa *Neoblattella*, *Cariblatta* and *Nahublattella*, but with the interesting addition of *Margattea,* which is a genus widely distributed in Asia, Oceania and Africa. *Margattea,* at least superficially, shares morphological characteristics with its Neotropical cousins (i.e., buffy body coloration, spotted pronotum). The next major clade was only supported in C100_Full (not the 2$^{nd}$ codon position or reduced data trees), *Supella* + *Sundablatta* + *Baltini*. The monophyly of Baltini was strongly supported, and here it constitutes *Balta*, *Ellipsidion*, *Pachnepteryx*, and *Saltoblattella*. *Ellipsidion* was previously thought to have a close relationship to *Balta (Roth 1999; Bourguignon et al. 2018)*. To our knowledge, no hypothesis about the relationship of *Pachnepteryx* was previously given, other than its assignment to Pseudophyllodromiinae (Roth 1998). The phylogenetic position of *Saltoblattella* has previously been debated. Bohn et al. (2010) discussed morphological evidence suggesting a relationship with Pseudophyllodromiinae, Ectobiinae or Blattellinae, with the latter position settled on only tenuously. Djernæs et al. (2012) recovered *Saltoblattella* with *Ectobius*, but with only a small genetic sampling. Now we recover it in Pseudophyllodromiinae with strong support as sister to an unidentified species of *Ellipsidion*. The final major clade of Pseudophyllodromiinae we term the Chorisoneurini. This clade includes only Neotropical genera: *Riatia*, *Anisopygia*, *Euthlastoblatta*, *Calhypnorna, Macrophyllodromia, Dendroblatta,* and *Chorisoneura.* Morphologically, this clade is quite diverse, and a few of these taxa were previously placed in another subfamily. Anisyutkin (2008) noted the remarkable similarity of *Anisopygia* to some *Ischnoptera*, which we agree with. However, Legendre et al. (2015) recovered, with strong support, *Anisopygia* with the Pseudophyllodromiine genus *Isoldaia* and which in turn was sister to *Dendroblatta* with low support (but possibly due to long branch attraction). Further testing is needed to safely determine the position of this enigmatic taxon. *Calhypnorna* is listed as Blattellinae on the Cockroach Species File online database (Beccaloni 2018) but this may be an error since Princis (1965) listed it in Anaplectidae and near *Plectoptera*, which would be considered Pseudophyllodromiinae by a modern concept. Evangelista et al. (2015) noted that previous genetic barcode analyses indicated a close relationship with *Chorisoneura* (Evangelista et al. 2013).

In Blattellinae we recover four principle clades. The first clade is a combination of Symplocini and Blattellini and nearly all relationships here are have maximal support. We recover *Episymploce sundaica* as a sister to *Blattella*, whereas Roth (1985) hypothesized it was more closely related to *Hemithyrsocera.* The second major clade is the Parcoblattini. This collection of genera (*Paratemnopteryx*, *Parcoblatta* and *Asiablatta*) was recovered with strong support in Bourguignon et al. (2018). Earlier, Roth (1990) proposed this clade on morphological grounds, although he did not examine *Asiablatta*. The next clade we recover is *Theganopteryx* + Hemithyrsocerini. *Theganopteryx* was previously thought to be in Ectobiinae, but Bohn et al. (2010) discussed why this was not supported. We give the name Hemithyrsocerini to the next clade, which is supported as including *Lobopterella*, and *Burchelia* by Roth (1988) and Rehn (1933) respectively. The last major clade of Blattellinae we recover is the Pseudomopini (also sometimes referred to as Ischnopterini or Ischnopterites). We recover *Ischnoptera* as close to *Euhypnorna*, which was thought to be close to *Calhypnorna* (Hebard 1921a) (and the superficially similar genera *Hypnorna* and *Hypnornoides*) but is actually more morphologically similar to *Ischnoptera* (Evangelista et al. 2019). We then find *Chorisoblatta* as sister to *Chromatonotus* (see next section). We recover *Beybienkoa* as sister to *Xestoblatta*. These genera have both, at various times (Hebard 1916; Legendre et al. 2015; Bourguignon et al. 2018), been proposed to be closely related to *Ischnoptera*, but this is the first time all three have been studied together. *Cahita*, *Dasyblatta* and *Pseudomops* were also recovered together, *Cahita* has been proposed as close to *Ischnoptera* (Rehn 1937) and *Dasyblatta* as close to *Chromatonotus* (Hebard 1921b). The three genera we recover together all have similarly hirsute bodies.

The relationships among the lineages of Blaberidae are not very well supported in our study. While we discuss some relationships here, we note that many of these are likely to change with new data and approaches.

A recent molecular studies did not support the monophyly of neotropical Epilamprinae (Legendre et al. 2017) but we recover this clade with moderate support. In accordance to recent molecular studies (Legendre et al. 2015; Legendre et al. 2017) we include *Thanatophyllum* in this group. However, the genus was originally placed in Zetoborinae (Grandcolas 1990).

In the C100_Full tree and most of the other trees inferred from 100 and 265 loci we recovered a clade we refer to as the Peri-Atlantic Blaberidae. All of the taxa constituting this clade (*Gyna,* Panchlorinae, *Aptera,* Blaberinae, and Zetoborinae) reside in Africa or the Neotropics. *Gyna* + Panchlorinae and *Aptera* + Blaberinae + Zetoborinae were both moderately supported in the C100_Full and C100_Reduced trees. More data on Gyninae will be needed to determine if this placement is strongly supported or not. A position close to Panchlorinae was recovered before (Evangelista et al. 2019) but other studies have other positions (Legendre et al. 2015; Bourguignon et al. 2018).

We recover a clade consisting of Blaberinae and Zetoborinae with moderate to strong support. The paraphyly of these taxa with respect to each other has been demonstrated previously (Legendre et al. 2017). The positions of most of the genera here disagree with previous phylogenies (Grandcolas 1998; Legendre et al. 2017; Bourguignon et al. 2018) so we refrain from discussing them until more data is collected.

The remaining Blaberidae fall into a weakly supported clade we call the Peri-Indian Blaberidae, since all of the taxa reside on land masses surrounding the Indian Ocean. Here, we recover Diplopterinae as sister to Oxyhaloinae, although support is weak. Diplopterinae has been a rogue taxon in past studies (Legendre et al. 2015; Legendre et al. 2017; Wang et al. 2017; Evangelista et al. 2018) but congruence over its position near Oxyhaloinae may be emerging (Bourguignon et al. 2018; Evangelista et al. 2019). We still remain cautious to accept any relationship without more evidence. Within Oxyhaloinae, we see *Oxyhaloa* as sister to all the other taxa, which has never been proposed before because no molecular study has included this taxon. We have the Gromphadorhini (Madagascar) nested within the African Oxyhaloinae, which has been proposed before (Legendre et al. 2017; Bourguignon et al. 2018). *Nauphoeta* and *Henschoutedenia* as sister group has strong support in our study and others (Legendre et al. 2015; Legendre et al. 2017). The remaining lineages of Blaberidae were all grouped together with little or no support. The only notable relationship we do recover with strong support is the monophyly of some old-world Epilamprinae, which has also been supported by other datasets (Legendre et al. 2017). More data, or improved modelling strategies, are necessary for further resolving these relationships.

#### Examining the position of *Chorisoblatta*

The position of *Chorisoblatta*, nested within a Neotropical group of Blattellinae is unusual given that this taxon is limited to Africa and Madagascar, and it is previously classified as a Pseudophyllodromiinae (Beccaloni 2018). Roth and Rivault (2002) hypothesized that there was a close relationship between *Chorisoblatta* and *Desmosia* Bolívar, 1895, who had similar morphology in their tarsal claws, femora, and (partially) male genitalia. It is possible that these two genera are members of Blattellinae with side reversed genital symmetry (with the hook on the right side, rather than the left). Yet, we are skeptical of this proposal because of the extreme similarity between the *Chorisoblatta* and *Chromatonotus* sequences. Of the 94 loci that contain data for both taxa only 19 contain any sequence differences, and only three loci contain more than four character differences. Therefore, the observed position of *Chorisoblatta* sp. could be due to cross contamination of samples. Future work will have to confirm this.

### Section 4: Other tests of locus quality

#### Confirming correction for trait correlation

Although we designed the test sets (Fig. 2, Table 1) to correct for correlation among locus traits (Fig. S1.1) it was not possible to omit all correlations entirely given the limited number of loci in our sample. Table 1 shows the correlation among traits in the test sets. To disentangle these effects, we did three more tests with a smaller number of loci for: mutation saturation, mean pairwise sequence distance (as inferred with the MG model), and dN/dS. For each, it was possible to effectively remove correlation with other factors by reducing the sample size down to 37 loci and allowing some of the treatments to overlap slightly (Fig. S4.1).

#### Figure S4.1

*Comparison of treatments for three scaled-down tests with extra trait correction. Boxes represent middle 75% quantiles; whiskers represent the remaining quantiles with points being outliers. White lines are the median value and black lines are the mean. N indicates how many loci are in each treatment for each factor.*

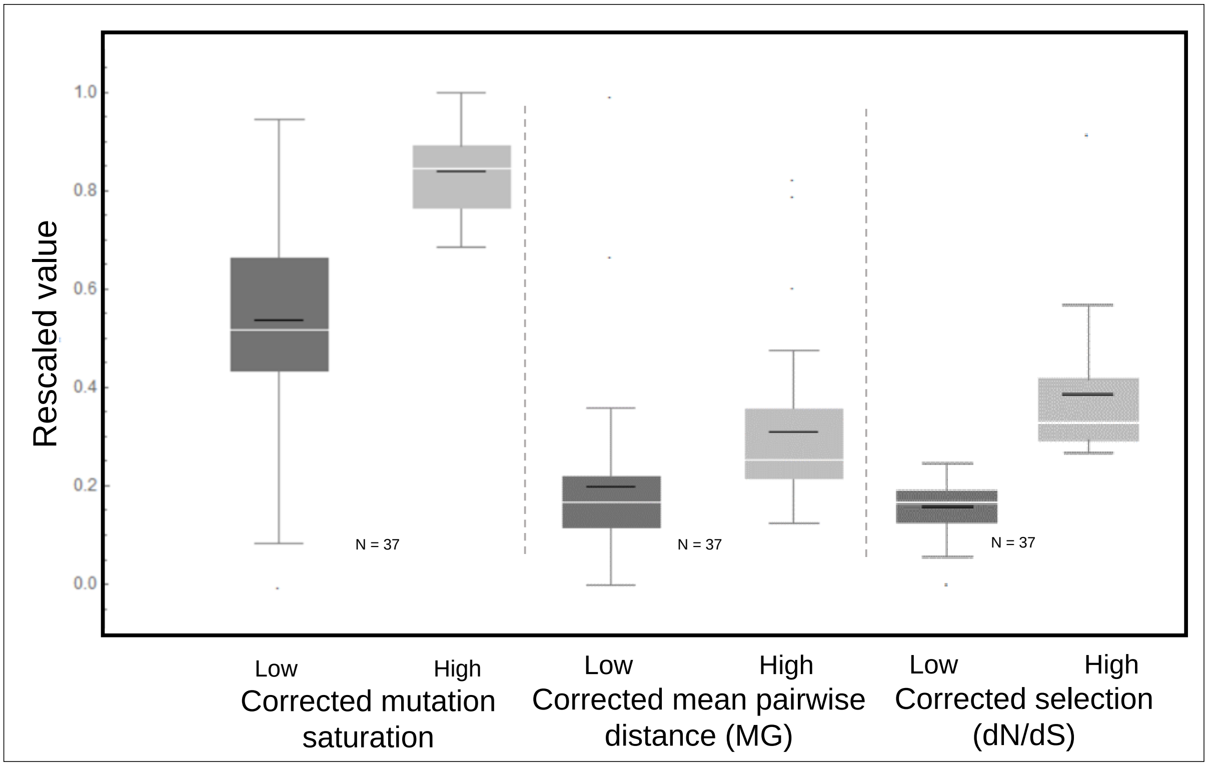

The results of the three scaled-down correction tests (Fig. S4.2) mirror that of the full tests (Fig. 4) except we see a stronger effect size here. In short, low mutation saturation, low mean pairwise sequence distance (MG model) and low selection are all traits that increase phylogenetic synecdoche with statistical significance.

We should note that, given our limited total number of loci, we could not design treatments that were independent from one another and also independent from locus length. Fortunately, the results that we found (Fig. S4.2) were not consistent with an interpretation based on locus length (i.e., the set with longer loci performed worse). Thus, the results of these tests are reliable with respect to locus length.

#### Figure S4.2

*Comparison of results for three scaled-down tests with extra trait correction. Plots show distribution of mean Robinson-Foulds distance to a set of baseline trees (inferred from a full dataset of 265 loci). Asterisks indicate statistically significant difference of distribution of means as determined by a Z-Test.*

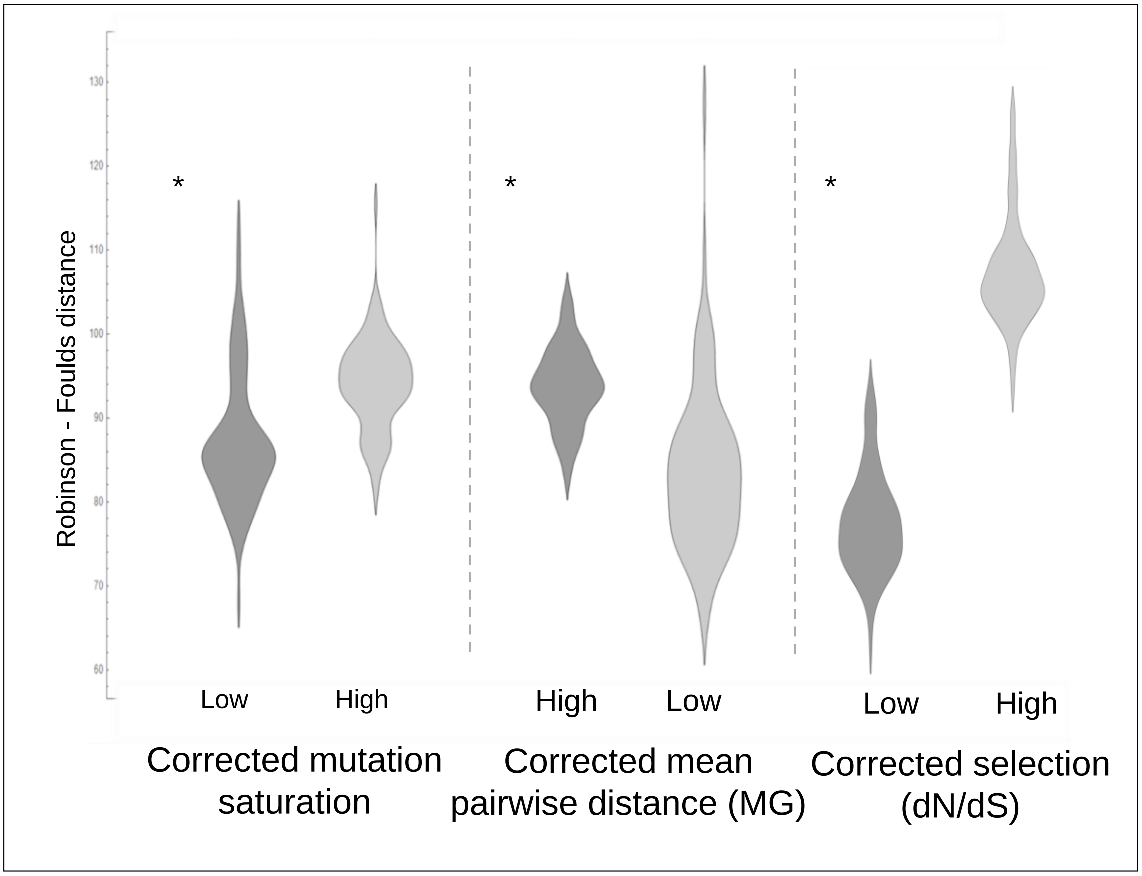

#### Tree precision via RADICAL

Here we present the results from the RADICAL experiments as comparisons of all trees at a given step across all iterations (i.e. not in comparison to a baseline as in Fig. 3). In other words, we show the effects of locus traits on tree precision. The results (Fig. S4.3), show that low mean pairwise sequence distance, fast mutation rate, and low information content visibly improved tree precision. Statistical comparison of these are shown in the main text (Fig. 4B).

##### Figure S4.3

*Changes in tree precision as measured by a RADICAL analysis. Points represent all Robinson-Foulds distances among trees at a given step for all iterations of RADICAL. The first step is omitted because it is not comparable with a linear model. We chose a linear model here, instead of an exponential model, primarily because values are expected to, and do indeed, reach 0, which is impossible under an exponential model.*

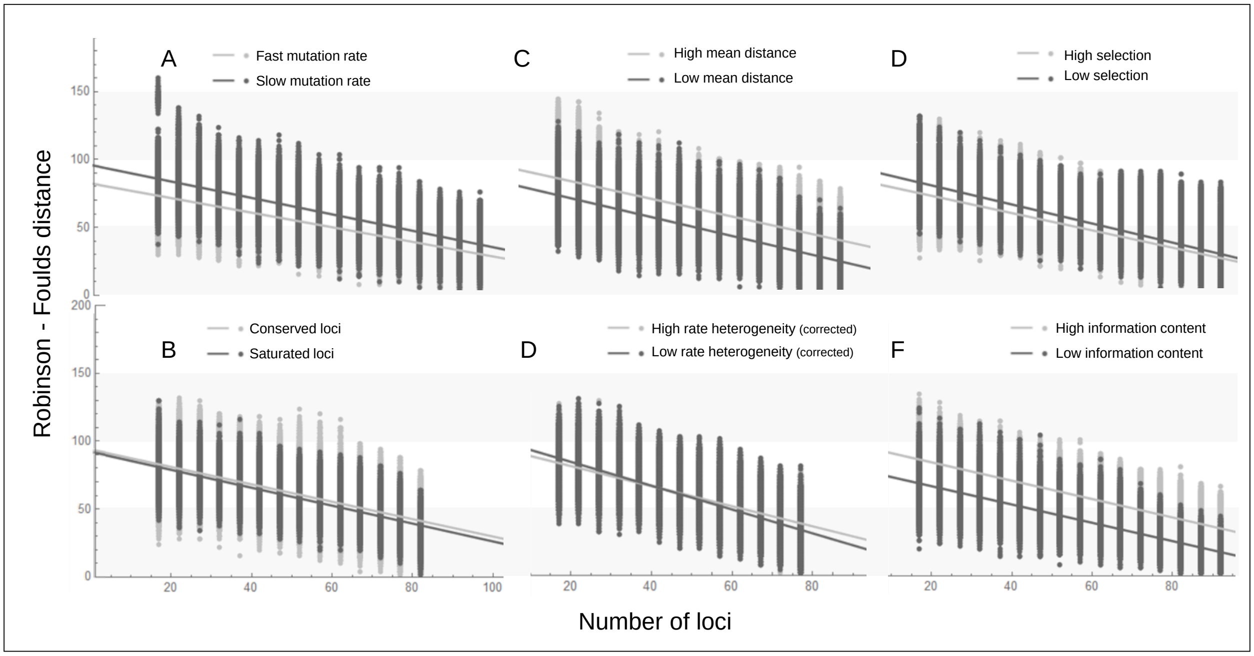

#### Other results

Although massive datasets may be detrimental to phylogenetic inference because of accumulated biases and noise, increases in the amount of overall data should be beneficial, particularly at the small scale.

To demonstrate this basic principle, we tested the effect of random concatenation of loci of different nucleotide lengths. We compared a set of 82 short loci (mean alignment length = 523.6) and 82 long loci (mean length = 1581.7; Fig. 2). These two sets were corrected for their number of taxa (means for sets 73.1 and 78.8 respectively), evolutionary rate (means 0.51 and 0.50 respectively), corrected rate heterogeneity (0.011, 0.009), and selection (0.049, 0.056; Table 1). Saturation (means 0.59 and 0.81 respectively) and mean pairwise sequence distance (2.0, 3.7) were correlated too strongly to correct for them (Table 1), but the results show trends opposite that of what would be expected for these two traits (Fig. S4.4).

To demonstrate the need to correct for locus length we tested the effect of raw rate heterogeneity, which is highly correlated with locus length. We compared a set of 87 low heterogeneity loci (mean number of rate categories = 5.7) and 87 high heterogeneity loci (mean rate categories = 13.7; Fig. 2). These two sets were corrected for their number of taxa (means for sets 79.0 and 82.8 respectively), evolutionary rate (means 0.49 and 0.49 respectively), and selection (0.054, 0.054; Table 1). As above, the correlation between mean pairwise sequence distance (2.24, 3.50) and saturation (0.66, 0.79) could not be corrected for (Table 1) but did not confound the results (Fig. S4.4).

As expected, datasets concatenating longer loci together showed drastically greater phylogenetic synecdoche (Fig. S4.4). Raw rate heterogeneity shows an unexpected trend, that higher heterogeneity improves phylogenetic inference (Fig. S4.4). We demonstrate that this is a result of the correlation with locus length because after controlling for it (Figs. 2, 3, 4, Table 1) the pattern is reversed and low rate heterogeneity is the more desirable feature.

#### Figure S4.4

*Results of RADICAL test of the effect of locus length and raw rate heterogeneity on phylogenetic synecdoche. Each point represents the mean Robinson-Foulds distance of a tree estimated in a RADICAL iteration to a set of baseline trees. Curves are exponential best-fit regressions.*

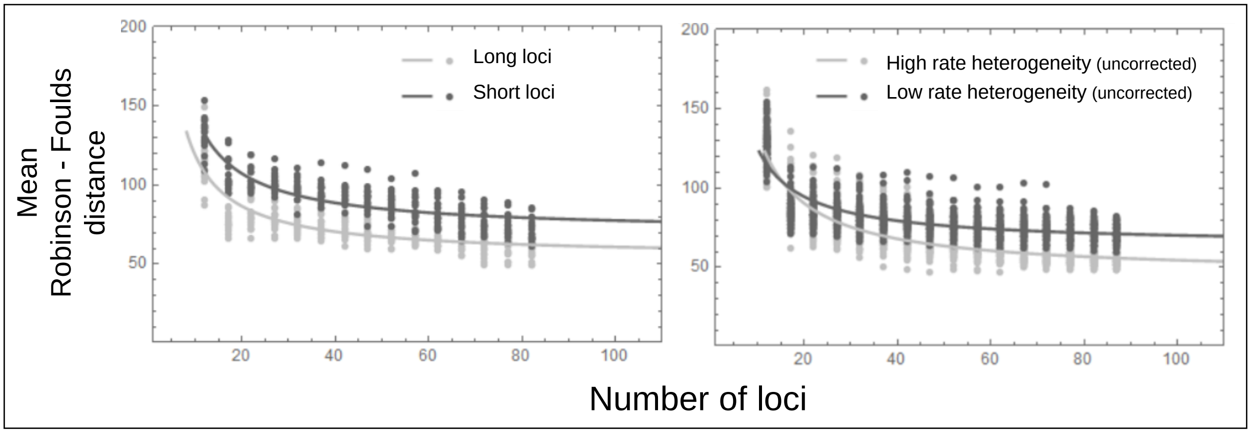

### Section 5: Evaluation of data reduction for phylogenetics

#### Topology from 265_Full and C100_Full

While there are many differences between the 265_Full and C100_Full phylogenies, their evolutionary scenario is largely equivalent (distance of 20 Robinson-Foulds units, when *Lanxoblatta* sp. is removed from the 265_Full tree; Fig. S5.1). The C100_Full tree was calculated as having a slightly lower ratio of external branch length to internal branch length (i.e. leafiness), one relative measure of how many characters were inferred to support relationships.

##### Figure S5.1

*Cophylogeny comparison of two phylogenetic hypotheses. Phylogenetic hypotheses correspond to the 265_Full and C100_Full analyses. Collapsed clades indicate equivalent split pattern between the two trees. Purple lines connect equivalent taxa.*

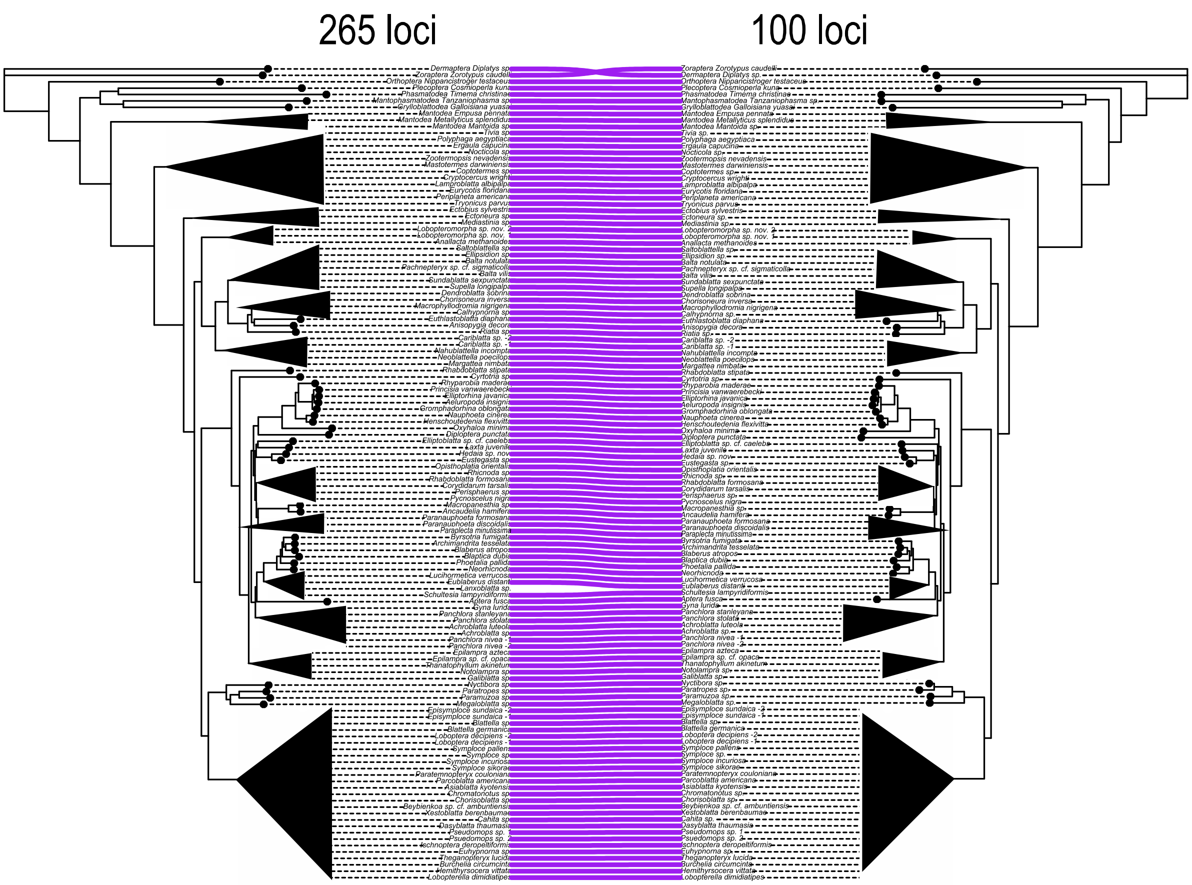

#### Tree quality and support

To assess phylogenetic similarity, the Robinson-Foulds distance was calculated between all main trees inferred in this study. The trees compared were those inferred from all 265 loci (265_Full), 100 of the best loci (not considering alignment taxon completeness; 100_Full), 100 of the best loci while considering taxon completeness (C100_Full), each alignment's 2$^{nd}$ codon positions and each alignment reduced to only those positions missing <50% of nucleotide data.

Table S5.1 shows the RF distances among all nine trees describe above. All trees inferred from only second nucleotides were very different from all other trees (most RF's >80), but when more data was included (i.e. 265_Full alignment) the 2$^{nd}$ nucleotide tree was more similar. The trees inferred from alignments that were reduced to the most complete nucleotide positions were most similar to each other but also moderately similar to the full alignments. Among the full alignments, the greatest similarity is seen from the trees C100_Full to 265_Full and from 100_Full to C100_Reduced.

The similarity seen among the C100_Full and 265_Full trees (RF = 20) demonstrate the high degree of phylogenetic synecdoche in the C100_Full alignment. The information content in this alignment is largely preserved, despite the loss of almost 2/3 of the data.

##### Table S5.1

*A symmetrical matrix of Robinson-Foulds distances among nine trees.*

| | 265_Full | 265_2nd | 265_Reduced | 100_Full | 100_2nd | 100_Reduced | C100_Full | C100_2nd | C100_Reduced |
|---|---|---|---|---|---|---|---|---|---|
| 265_Full | 0 | 60 | 56 | 48 | 98 | 66 | 20 | 96 | 44 |
| 265_2nd | 60 | 0 | 84 | 82 | 84 | 88 | 64 | 80 | 78 |
| 265_Reduced | 56 | 84 | 0 | 30 | 96 | 36 | 54 | 92 | 24 |
| 100_Full | 48 | 82 | 30 | 0 | 96 | 40 | 40 | 94 | 20 |
| 100_2nd | 98 | 84 | 96 | 96 | 0 | 104 | 96 | 92 | 104 |
| 100_Reduced | 66 | 88 | 36 | 40 | 104 | 0 | 60 | 92 | 28 |
| C100_Full | 20 | 64 | 54 | 40 | 96 | 60 | 0 | 90 | 44 |
| C100_2nd | 96 | 80 | 92 | 94 | 92 | 92 | 90 | 0 | 92 |
| C100_Reduced | 44 | 78 | 24 | 20 | 104 | 28 | 44 | 92 | 0 |

In addition to quantifying tree similarity, the plausibility of each tree was tested statistically using an approximately unbiased (AU) test. We tested the plausibility of the nine trees given the three full alignments (265_Full, 100_Full, C100_Full).

The trees deemed plausible (p-values > 0.05) were those inferred from 265_Full, 100_Full, C100_Full and C100_Reduced (Table S5.2). Each of the "full" trees were plausible under the alignments they were inferred from, and the C100TFull was also plausible under the 100_Full alignment. The C100_Full was the only tree deemed plausible for more than one alignment.

Table S5.2

*P-values for nine trees given three alignments in an AU-test*

| | | P-value given alignment in AU test | | | | | |
|---|---|---|---|---|---|---|---|
| | | 265_Full alignment | | 100_Full alignment | | C100_Full alignment | |
| Trees | 265_Full | 0.9987 | * | 0.0311 | - | 0.0226 | - |
| | 265_2nd | 0.0002 | - | 0 | - | 0.0013 | - |
| | 265_Reduced | 0 | - | 0.0059 | - | 0.0006 | - |
| | 100_Full | 0.0001 | - | 0.8492 | * | 0.0001 | - |
| | 100_2nd | 0 | - | 0 | - | 0 | - |
| | 100_Reduced | 0 | - | 0.0158 | - | 0 | - |
| | C100_Full | 0.0013 | - | 0.1541 | * | 0.9802 | * |
| | C100_2nd | 0.0006 | - | 0 | - | 0 | - |
| | C100_Reduced | 0 | - | 0.3606 | * | 0.0014 | - |

[a] "-" indicates statistically significant rejection of the tree given the alignment.

[b] "*" indicates that the tree could not be rejected and was plausible given the alignment

The p-values from the AU tests provide a means of accepting or rejecting the trees based on distribution of lnL's from 10,000 RELL bootstraps. Yet the lnL's, or the difference in lnL from the best (ΔlnL), can also be used to rank the trees.

Table S5.3 gives the ΔlnL values for each of the nine trees tested under all three alignments. Under the 265_Full alignment, the most plausible evolutionary scenario was that of the 265_Full tree. Ranked second under the 265_Full alignment is the C100_Full tree. For the 100_Full alignment, the C100_Reduced and then 265_Full trees were second and third most likely and for the C100_Full alignment 265_Full was second most likely. Thus, C100_Full was ranked highly under each alignment.

#### Table S5.3

*Delta log-likelihood (ΔlnL) values for nine trees under three alignments as determined by the approximately unbiased (AU) test.*

| | | **ΔL given alignment** | | |
|---|---|---|---|---|
| | | 265_Full alignment | 100_Full alignment | C100_Full alignment |
| **Trees** | 265_Full | 0 | 87.229 | 64.107 |
| | 265_2nd | 1609.873 | 1090.407 | 1034.373 |
| | 265_Reduced | 506.154 | 117.254 | 356.327 |
| | 100_Full | 464.938 | 0 | 202.124 |
| | 100_2nd | 3295.582 | 1005.028 | 1672.214 |
| | 100_Reduced | 839.734 | 94.035 | 430.82 |
| | C100_Full | 131.413 | 51.709 | 0 |
| | C100_2nd | 3572.893 | 1320.567 | 1508.716 |
| | C100_Reduced | 432.121 | 17.669 | 245.832 |

Thus, we show above and in the main text that the C100_Full tree displays a high degree of phylogenetic synecdoche and has a number of quantifiable (e.g., decreased leafiness, high support under the AU test) and subjective (e.g., inferred from less noisy data) improvements. Despite this, the bootstrap and internode certainty scores are marginally lower in C100_Full to 265_Full, potentially leading one to the conclusion that the reduction of the dataset has decreased support for the tree. Perhaps, this is the case. However, reducing the number of characters in any alignment will affect node support values, regardless of the information content of the characters removed (Soltis and Soltis 2003; Klötzl and Haubold 2016). Indeed, phylogenies inferred from extremely long alignments typically have very high bootstrap support values (e.g.,Misof et al. 2014; Garrison et al. 2016; Evangelista et al. 2019), but that support can be highly dependent on only a small number of nucleotides (Shen et al. 2017; Simon et al. 2018). As alignment lengths increase towards infinity, a bootstrap subsample of that alignment would be identical to the original alignment it is drawn from and thus 100% support will be found with certainty (Soltis and Soltis 2003). Reducing the length of the alignment may allow bootstrap values to fall in range that is informative.

### Section 6: Hands-on guide to improving phylogenies through optimized subsampling

#### Part A: De novo assessment of subsample optimality

*Note: This part is specifically for those who want to assess which loci are optimal using their own predefined criteria. Start at Part B if the relationship between locus features and locus quality has already been established.*

**Step A-1:** Using a multi-locus dataset, infer three or more "full data" phylogenies from as many loci as possible. The trees should be inferred with different methods to account for error in tree inference. For example, in this study we inferred three trees: one with all the data, one with only second nucleotides (to lessen error from mutation saturation), and one alignment with only the most taxon-complete nucleotide positions (to lessen error from missing data patterns).

*Note: If your goal in generating optimized subsamples is to reduce the costs of data collection you will need a preliminary dataset, likely from a reduced sample of taxa. However, your goal may be to improve computational efficiency and tree quality using an existing phylogenomic dataset, as in our study.*

**Step A-2:** Identify locus qualities that may or may not improve phylogenetic inference. This may be determined by the assumptions or limitations of the evolutionary models you wish to use. Our approach examined some predefined locus qualities (e.g., mutation rate, rate heterogeneity, saturation, mean pairwise sequence distance, selection). Another approach would be to use loci identified by a software like PhyInformR (Dornburg et al. 2016), although we personally prefer tree independent methods.

**Step A-3:** Design a controlled experiment to test the quality of each phylogenetic subsample under the criterion of phylogenetic synecdoche. We used the RADICAL subsampling protocol (Narechania et al. 2012) to compare the rates and extent to which each locus set converged on the "full data" tree topologies. However, other methods of comparison (e.g., single tree inference from each subsample, locus jack-knifing, locus bootstrapping) could be appropriate too. The only requirement to use the criterion of phylogenetic synecdoche is to compare tree(s) inferred from each subsample against the trees inferred from the full dataset.

**Step A-4:** Assess the experimental results. The subsample whose tree (or trees) has more similarity to the full-data trees (i.e. demonstrate phylogenetic synecdoche) should be treated as more optimal. We estimated tree similarity using the Robinson-Foulds distance metric (Robinson and Foulds 1981) but a composite distance metric is likely more meaningful (Kuhner and Yamato 2015), particularly if your dataset contains rogue taxa or your experimental design did not utilize high numbers of replicates. A valid statistical comparison of tree distances should be made depending upon the nature of your experiment.

#### Part B: Composing optimized subsamples and inferring an improved phylogeny.

**Step B-1:** Assign scores to loci based on optimality criteria. A simple way of composing optimized subsets is to rescale locus traits based on which polarity of the trait is more optimal. For example, if low rate heterogeneity is a desired characteristic, rescale all loci rate heterogeneities to values from 0-1 with 1 being the lowest heterogeneity and 0 being the highest heterogeneity. Do this for all locus traits so that 1 is assigned to the more optimal trait polarity and 0 is assigned to the least optimal trait polarity. See note below for an optional step here. Now sum all the rescaled traits to assign a locus an optimality score and do this for all loci.

*NOTE: Another step that can be added here is to weight the rescaled values. For instance, we found that loci with low mean pairwise sequence distance performed far better than loci with high pairwise distance. In contrast, we found that loci with low selective pressure were only slightly better than loci under high pressure (perhaps because our loci were all under strong stabilizing selection). Therefore, it would be justifiable to up-weight the rescaled mean pairwise sequence distance values, possibly doing so relative to the traits effect size.*

**Step B-2:** Choose the most optimal loci. Choose the loci with the highest overall score. The number of loci one should choose is determined by the number of taxa in the tree, the density of information content per locus, the amount of data completeness, the computational power available and the types of further data reduction analyses desired. These are all up for debate and many are the preference of the investigator. We arbitrarily chose to reduce to 100 loci (66% reduction), which was sufficient for recovering a robust tree efficiently but not sufficient to yield meaningful support values from further data reduction analyses.
